# Cell free expression in proteinosomes prepared from native protein-PNIPAAm conjugates

**DOI:** 10.1101/2023.08.23.554401

**Authors:** Mengfei Gao, Dishi Wang, Weihua Leng, Michaela Wilsch-Bräuninger, Jonathan Schulte, Nina Morgner, Dietmar Appelhans, T-Y. Dora Tang

## Abstract

Towards the goal of building synthetic cells from the bottom-up, the establishment of micrometer-sized compartments that contain and support cell free transcription and translation that couple cellular structure to function is of critical importance. Proteinosomes, formed from crosslinked cationized protein-polymer conjugates offer a promising solution to membrane-bound compartmentalisation with an open, semi-permeable membrane. Critically, to date, there have been no demonstration of cell free transcription and translation within water-in-water proteinosomes. Herein, we present a novel approach to the fabrication of proteinosomes directly from native protein-polymer (BSA-PNIPAAm) conjugates. We show that these native proteinosomes offer an excellent alternative as artificial cell chassis. Significantly, the native proteinosomes are stable under high salt conditions and can consequently support cell free transcription and translation. The native proteinosomes offer enhanced protein expression compared to proteinosomes prepared from traditional methodologies. Furthermore, we demonstrate the integration of proteinosomes into higher order cellular architectures with membrane free compartments and liposomes. The integration of bioinspired architectural elements with the central dogma is an essential building block for realizing minimal synthetic cells and is key for exploiting artificial cells in real-world applications.

## Introduction

A grand challenge in synthetic biology is the bottom-up construction of synthetic cells that have life-like properties. The integration of bioinspired architectural elements with enzymatic function is a minimal requirement for the generation of synthetic cells. Thus, the facile establishment of micrometer-sized compartments which contain and support cell free expression systems provide the foundation to build robust synthetic cells with increased functional and structural complexity. A prime example has been the incorporation of cell free expression systems into lipid vesicles ^[1,2]^ that have been pivotal in realizing cellular structure and function in a minimal fashion and has been an essential foundational technology in synthetic cellular research. This system provides a lipid bilayer boundary that contains cell free expression system that models the biological cell with its cell membrane and internalized transcription and translation. This platform has not only been used to model noise ^[3]^ and crowding in the biological cell ^[4]^ but has also been used as prototype to incorporate higher order cellular functions such as lipid synthesis^[5,6]^ DNA replication^[7]^ and protein induced membrane deformations.^[8]^

Notwithstanding lipid vesicles, there have been a variety of different compartments which can support cell free expression from water-in-water membrane bound compartments such as polymersomes ^[12,13]^ to water-in-oil systems such as surfactant stabilized droplets and Droplet Interface Bilayers (DIBS),^[14]^ inorganic colloidosomes,^[15]^ and membrane free systems such as coacervates ^[16]^ and hydrogel-based systems ^[23]^. These synthetic cellular platforms have been important for increasing compartment stability *via* polymer stabilization ^[17]^ for enhancing properties such as intercellular communication and for integrating higher order cellular architectures ^[18–20]^ that can be critical for integrating specific cellular properties into non-living matter.

Despite the growing number of ways to include cell free expression systems into water-in water membrane bound compartments there remain some challenges with their utilization as synthetic cellular platforms. Namely, that aqueous membrane-bound compartments, liposomes and polymerosomes are closed systems which contain a finite amount of resources. This limits the length of time that expression can take place and the long-term function of the compartment. The addition of membrane pores ^[9,10]^ or light activated lipids^[11]^ can tune membrane permeability to external substrates thus providing more resources to internalized cell free expression systems. The establishment of these systems are non-trivial and every additional component increases the molecular complexity of the system and increases the challenges in their fabrication. Therefore, there is still a necessity for alternative solutions for membrane bound compartmentalized cell free expression systems that provide new functions and features to increase the flexibility and functionality of synthetic cells. Expanding the repertoire of synthetic cellular platforms provides the ability to open and extend the range of applications and functionalities by selecting specific properties as an end user. To this end, proteinosomes are alternative membrane-bound compartments prepared from protein-polymer conjugates^[19,20]^ with membranes that are elastic with tunable permeability to enzymes or DNA molecules.^[21–24]^ Water-in-water proteinosomes have been shown to support reaction cascades,^[25]^ DNA strand displacement reactions and PEN DNA reactions.^[20,21]^ They can be manipulated to generate multi-compartments systems^[24,26–28]^; and are compatible with natural cells,^[29]^ responsive to stimuli,^[30]^ can be chemically tuned to incorporate light activatable sorting communities^[31]^ and produced in a high throughput manner.^[32]^ Whilst cell free expression has been demonstrated within non-crosslinked BSA-PNIPAAm conjugates in oil^[25]^, there is no example of the cell free expression system within water-in-water proteinosomes. Integration of cell free expression within water-in-water proteinosomes provides a viable alternative for a synthetic cellular platform, an open compartment with a permeable membrane to allow the continuous supply of nutrients to the reaction site.

As described previously, the novelty of production of the proteinosomes lies in the generation of amphiphilic nanoparticles produced by conjugated protein with poly-(N-isopropylacrylamide) PNIPAAm. These protein-polymer conjugates provide the scaffold for the proteinosome by stabilizing water-oil emulsions which are then covalently crosslinked within the emulsions. Typically, protein-(PNIPAAm) conjugates are synthesized by cationization of a protein such as bovine serum album (BSA) *via* carbodiimide chemistry to increase the number of amine groups on the protein. The amine groups are used to conjugate end capped mercaptothiazoline-activate PNIPAAm to the protein and to provide crosslinking sites between the amphipathic protein-PNIPAAm conjugate to stabilize water-in-oil emulsions. To this end, pegylated bis(sulfosuccinimidyl)suberate (BS(PEG)_n_ crosslinks the protein-PNIPAAm conjugate via covalent links between the primary amine on the protein and the BS group. This provides the structural stability to the membrane. Furthermore, (BS(PEG)_n_ is commercially available with different PEG lengths which tunes the pore size and permeability of the membrane ^[23]^. Removal of the oil phase and transfer to water leaves water-in-water crosslinked protein-PNIPAAm compartments.^[32]^

Our primary goal was to achieve cell free transcription and translation within proteinsomes, however, when we produced proteinosomes using the conventional methods using cationized protein-polymer conjugates we found that the proteinosomes were not stable in the cell free expression systems, most likely due to the high salt content of the buffer. The stability could be limited by the protein cationization step in the preparation of the proteinosome where 1-(3-Dimethylaminopropyl)-3-ethylcarbodiimide hydrochloride (EDC) reactions are used to chemically modify the protein to add primary amine groups to aspartic and glutamic acid groups of the protein. Unfortunately, this process has been shown to denature enzymes which could affect the stability of the proteinosome.

Therefore, to integrate cell free expression systems with proteinosomes, we removed the cationization step to provide a simplified methodology for proteinosome preparation. We show that this simplified procedure provides robust proteinosomes. We exploited the porous membrane of the proteinosome to isolate plasmid DNA and allow transcription and translation machinery to diffuse across the membrane and drive transcription and translation within the interior of the proteinosome (See supplementary methods). We show that DNA and mRNA are retained within the proteinosome whilst expressed proteins diffuse out of the interior of the proteinosomes. The ability to encapsulate cell free transcription and translation within proteinosome will give added value to a synthetic cell community by offering an alternative synthetic cell chassis with facile methods of formation and an open membrane which can allow the diffusion of resources to its core. This provides a new platform for establishing higher order architectural structures, reactions and gene circuits with potential applications in living technologies.

## Results and Discussions

Given that cationized proteinosomes were not stable in the high salt buffer used for cell free expression. We developed a new methodology for the preparation of proteinosomes to remove the protein cationization step and reduce the excess positive charges that could hinder cell free transcription and translation (**Figure 1A**). To do this, we directly conjugated PNIPAAm capped with a mercaptothiazoline terminal group (**Figure S1, S2**) to the -NH_2_ group in lysines on the protein BSA that had not been subjected to cationization. The native BSA conjugate (nat-BSA-PNIPAAm), was characterized using Laser Induced Liquid Beam Ion Desorption (LILBID) mass spectrometry (**Figure 1B**) and Dynamic Light Scattering (DLS) to confirm that the polymer had been conjugated to protein. DLS data showed a difference in size distribution of the protein conjugate at 20 °C compared to 37 °C for the nat-BSA-PNIPAAm that was not observed in BSA alone (**Figure S3**). This correlated to the aggregation of conjugated protein-PNIPAAm conjugates above the melting temperature of the PNIPAAm chain as previously shown.^[33]^ In addition, mass spectrometry data showed peaks corresponding to BSA (67 kDa) and with peaks at 75, 83, 92 and 101 kDa which correlate to the molecular weight of nat-BSA-(PNIPAAm)_1_, nat-BSA-(PNIPAAm)_2_, nat-BSA-(PNIPAAm)_3_, nat-BSA-(PNIPAAm)_4_ respectively (**Figure 1B**). Together this data confirmed that PNIPAAm was conjugated to the protein and the average molecular weight of the PNIPAAm chain was 8.4 kDa. Given the molecular weight of PNIPAAm by mass spectrometry, we determined the average molar ratio of BSA: PNIPAAm was 1:5.7 by separately measuring the concentration of protein by a protein bicinchoninic acid assay and UV spectroscopy and PNIPAAm by UV absorbance at 300 nm (see materials and methods, **Figure S4 Table S1**).

**Figure 1:**
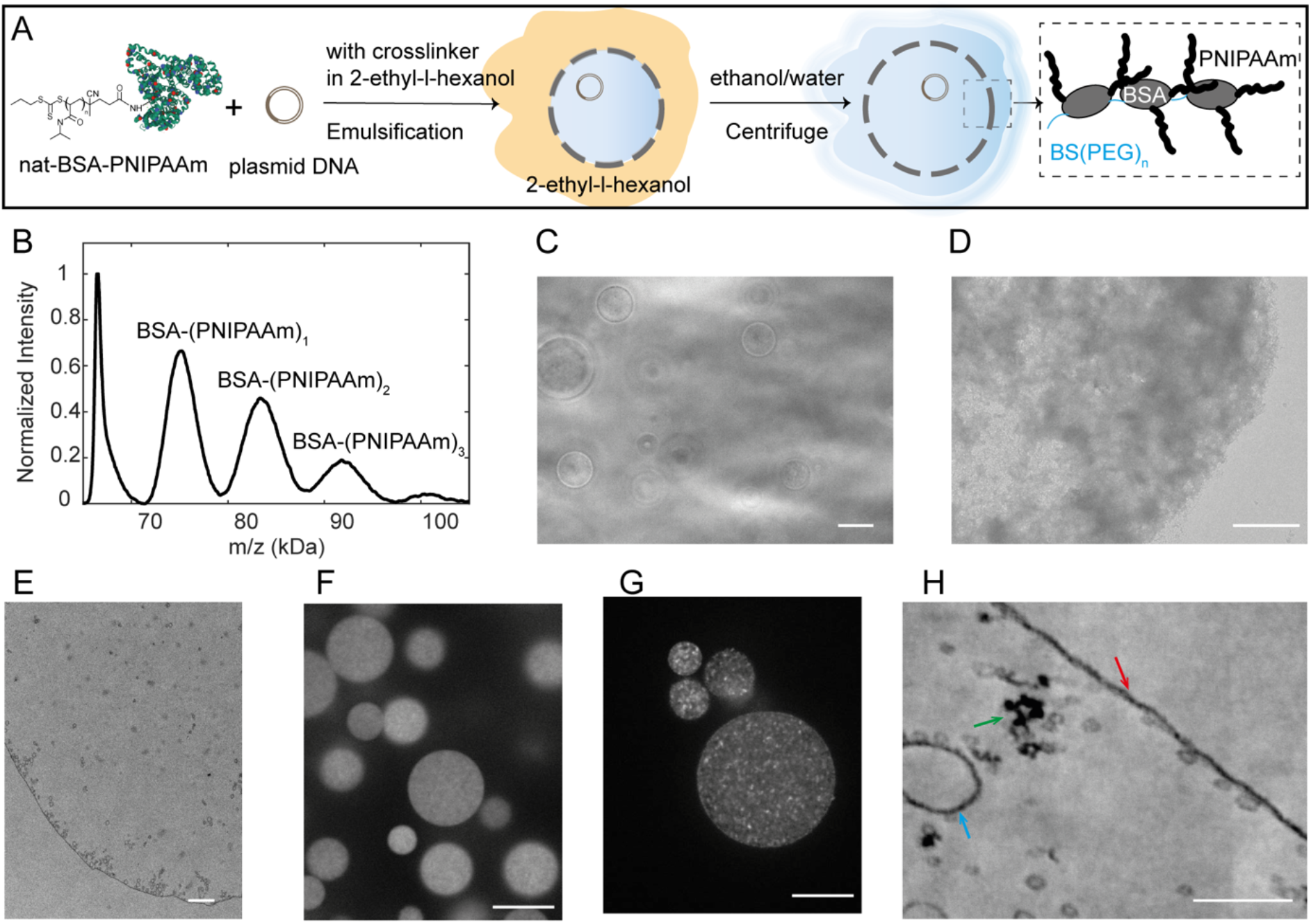
Preparation and characterization of native BSA proteinosomes. A) Schematic describing the preparation of the native proteinosomes. Nat-BSA-PNIPAAm and plasmid DNA are mixed in 0.1 M sodium bicarbonate (NaHCO_3_, pH 8.4) aqueous phase and mixed with 2-ethyl-l-hexanol containing BS(PEG)_n_ generating emulsions. After crosslinking, step-wisely purified from oil to water. B) Laser induced liquid beam ion desorption (LILBID) mass-spectra of nat-BSA-PNIPAAm. The average distance between multiple BSA-(PNIPAAm)_n_ peaks was calculated to be 8.4 kDa. C) Phase-contrast microscopy image of empty nat-BSA proteinosomes. Scale bar: 20 µm. D) Transmission electron microscope (TEM) image of a part of an empty nat-BSA proteinosome negatively stained with 1% uranyl acetate. Scale bar: 1 µm. E) TEM image of part of an empty nat-BSA proteinosome embedded in resin and cross-sectioned to a thickness of 70 µm. Scale bar: 1 µm. TEM images were obtained on a Tecnai 12 TEM. F) Fluorescence microscopy images showing plasmid DNA encapsulated within nat-BSA proteinosomes. G) Fluorescence microscopy images showing plasmid DNA encapsulated within BSA-NH_2_ proteinosome. Proteinosomes in F and G were stained with DNA dye EvaGreen and the images were obtained using a spinning disk microscope with a 60x/1.3 NA UPLSAPO objective. Scale bars in E and F: 10 µm. H) TEM image of a plasmid DNA contained nat-BSA proteinosome incubated with diluted PURExpress. The proteinosome sample was embedded in resin, cross-sectioned to a thickness of 70 µm and imaged on a Delong TEM at 25 kV. Red arrows refer to the proteinosome membrane, green arrow points to plasmid DNA, blue arrow points to small compartments. Scale bar: 100 nm. The images shown are representations of at least three experimental replicates. The replicates can be found in the data archive.

Having shown that end-capped mercaptothiazoline-activated PNIPAAm can directly conjugate to native BSA at a molar ratio of 1:5.7 (BSA:PNIPAAm), we tested the capability of nat-BSA-PNIPAAm conjugates to be coupled with PEG_2000_ *via* a bis(N-succinimidly succinate) group (BS(PEG)_2000_) to generate crosslinked membrane bound compartments. Nat-BSA-PNIPAAm was diluted in sodium bicarbonate solution and mixed with 2-ethyl-1-hexanol with BS(PEG)_2000_ at a volume ratio (φ) of 0.06. Water-in-oil emulsions were generated by pipetting the mixture 10 times at constant speed, the dispersion was then left overnight at 8 °C for crosslinking (see materials and methods). The crosslinked proteinosomes were transferred to water by centrifuging the dispersion at 3000 rcf and replacing the aqueous volume with 70% ethanol water, followed by repeated centrifugation and replacement of the aqueous volume with 50% and then 25% ethanol and water.

Optical microscopy images showed spherical proteinosomes indicating that the native protein polymer conjugates could be crosslinked and were stable in water (**Figure 1C**). Electron microscopy images obtained from negative staining of the proteinosomes confirmed that the proteinosomes were globular spheres delineated by a membrane (**Figure 1D**). Further characterization of proteinosomes embedded in resin, microtomed and imaged by electron microscopy showed a membrane of approximately 10-20 nm encapsulating a lumen (**Figure 1E**). Given the resolution of the imaging, the membrane thickness is in the range of single or double protein layers where the hydrodynamic diameter of BSA is approximately 4 nm. The slightly higher dimension could be caused by osmium tetroxide “staining” – which will react with the double-bonds within the proteins and slightly increase the thickness of the membrane layer. Transmission electron microscopy images also showed presence of small regions of increased electron density with a diameter of hundreds of nanometers. These small structures could be attributed to small crosslinked inverted micelles of protein-PNIPAAm conjugates that form during the preparation stage.

Given that stable proteinosomes could be formed from nat-BSA-PNIPAAm, we next tested the capability of these proteinosomes to support cell free transcription and translation. To do this, we chose to utilize the porosity of the proteinosome membrane to contain plasmid DNA and allow the diffusion of the transcription and translation machinery through the membrane. Therefore, we directly encapsulated circular plasmid DNA (2.3 MDa) into the proteinosomes by including the plasmid DNA into the aqueous solution prior to mixing with oil and preparation of emulsions by pipetting (see materials and methods). After transfer of the proteinosomes into water, EvaGreen dye (final concentration 10 µM) was added to the proteinosomes to image the DNA. Optical microscopy images showed DNA distributed within the interior of the proteinosomes (**Figure 1F**). Electron microscopy images of nat-BSA proteinosomes encapsulating DNA showed small regions of increased electron density attributed to plasmid DNA which were not present in the empty proteinosomes (**Figure 1H, green arrow**). Together, these results confirm that DNA could be encapsulated within the nat-BSA proteinosomes. Comparisons of optical microscopy images with proteinosomes encapsulating DNA prepared from cationized BSA (BSA-NH_2_) proteinosomes (**Figure 1G, Figure S5**) showed regions of high fluorescence intensity which could be attributed to aggregate formation; a lower encapsulation efficiency and a larger average size of proteinosome compared to the nat-BSA proteinosomes (**Figure 1F, Figure S6**). It could be possible that DNA interacts with the BSA-NH_2_-PNIPAAm conjugate which could affect conjugate-conjugate interactions and thus the physical properties of the proteinosomes.

Having confirmed that plasmid DNA could be encapsulated within the native proteinosomes, we then tested the ability for the native proteinosome to support cell free transcription and translation. To do this, we added nat-BSA proteinosomes that contained plasmid DNA to PURExpress and monitored transcription of mRNA and translation of protein in 4 µL of sample using a microplate reader (**Figure 2A**). DFHBI dye was added to the PURExpress to monitor mRNA production *via* a Broccoli RNA aptamer and protein production was detected *via* the expression of the red fluorescent protein, mCherry. Our results showed fluorescence increase from mRNA and protein production (**Figure 2B**). Furthermore, optical microscopy images showed that mRNA was localized within the proteinosome with increasing fluorescence intensity from mCherry from 10 to 170 min which is evenly distributed throughout the dispersion (**Figure 2C**). In order to confirm mCherry expression at low concentrations of proteinosomes, RFP trap beads were added to the solution on the exterior of the proteinosomes to capture mCherry protein (**Figure 2D, Movie S1**). Fluorescence microscopy imaging showed increased DHFBI signal inside the proteinosome (**Figure 2E**) and mCherry fluorescence intensity on the RFP trap over time (**Figure 2F**). These results show that native proteinosomes that encapsulate plasmid can support transcription and translation.

**Figure 2:**
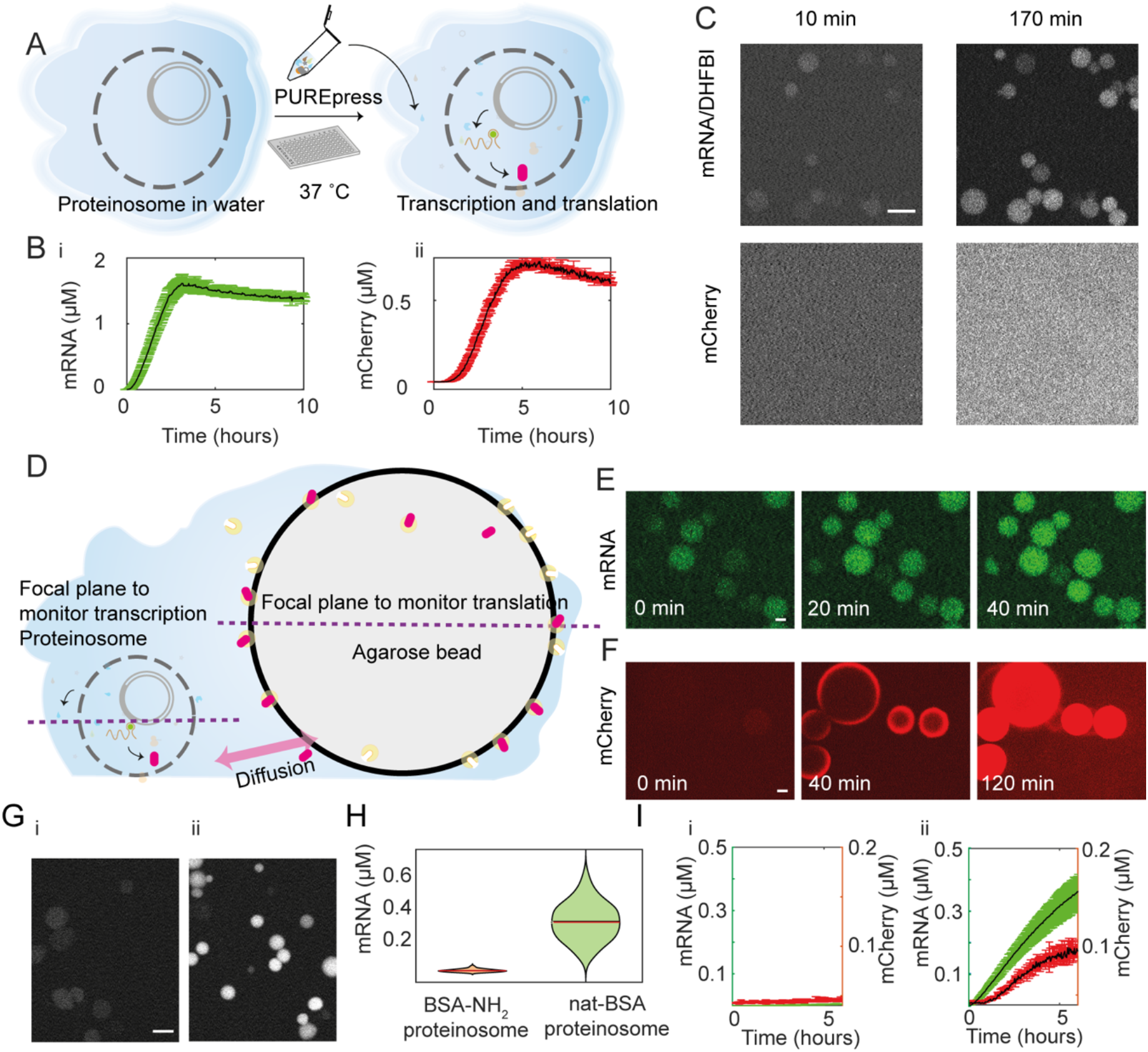
Cell free expression within BSA proteinosomes A) Schematic of experimental work flow for cell free expression within proteinosomes. Proteinosome in aqueous phase were mixed with PURExpress and imaged on a microplate reader. B) The kinetics of transcription of mRNA (i) and translation of mCherry (ii) were measured. C) Confocal microscopy images of nat-BSA proteinosomes containing plasmid DNA incubated with PURExpress after 10 mins and 170 mins. Scale bar: 10 µm. D) Schematic showing the incorporation of RFP trap with proteinosomes to confirm protein expression. Proteinosomes and the agarose beads-based RFP trap were imaged with confocal microscopy at different focal planes. E) Confocal microscopy images of DNA transcription in a dispersion of proteinosomes. F) The corresponding mCherry signals from the agarose beads in the same field of view but at different focal plane. G) Confocal microscopy images showing mRNA production in BSA-NH_2_ proteinosome (i) and nat-BSA proteinosome (ii) with PURExpress (0.33x) after 3 hours. H) The DHFBI/mRNA signal distribution extracted from G. Red line represents the mean value. I) Cell free expression in BSA-NH_2_ proteinosome (i) or nat-BSA proteinosome with PURExpress (0.33x) (ii), where transcription (green) and translation (red) were monitored with a Tecan Spark 20M microplate reader. Scale bar: 10 µm. All experiments were undertaken at 37 °C. 3 repeats were performed and the shade were plotted as mean (black line) ±standard deviation. All microscopy images are representations of the results obtained from replicate experiments.

To confirm that protein expression was taking place within the proteinosome, we undertook two control experiments. First, we gently centrifuged the proteinosomes to the bottom of an Eppendorf tube and took out the supernatant. The supernatant was added to PURExpress (1x concentration) and transcription and translation was monitored using a microplate reader (**Figure S7**). The results showed no fluorescence intensity from production of mRNA or protein which is commensurate with no transcription and translation taking place within the supernatant and confirming that all the DNA was contained within the interior of the proteinosomes. Secondly, plasmid DNA (2.7 nM) and PURExpress (1x) was gently mixed with empty nat-BSA proteinosomes and incubated for 3 hours at 37 °C. Confocal imaging of the dispersions incubated with either EvaGreen dye or DFHBI showed green fluorescence on the exterior of the proteinosomes with no fluorescence in the interior of the proteinosomes. This indicates that DNA and mRNA cannot diffuse through the membrane (**Figure S8**) and in the case where the DNA has been pre-encapsulated within the proteinosomes the results confirm that both transcription and translation take place within the proteinosome. Taking into account our earlier results, this shows that DNA and mRNA are retained within the proteinosome and the protein can diffuse through the membrane. This feature could be exploited in building communication channels *via* protein diffusion between proteinosomes.

When comparing the ability for BSA-NH_2_ proteinosomes to support transcription and translation to the nat-BSA proteinosomes, we found that BSA-NH_2_ proteinosomes disassembled in 1x PURExpress (**Figure S9**). It is possible that the collapse of the proteinosome was driven by the high salt content either by driving polymer collapse or due to specific interactions between the components within PURExpress and the proteinosomes. An alternative explanation could be attributed to the different elasticity of the membrane in native and cationized proteinosome that could arise from different degrees of crosslinking. As the cationized proteinosome have additional NH_2_ groups compared to native protein-polymer conjugates, there are more sites for crosslinking that could lead to membranes which are less porous and more rigid. The addition of PURE (1x) to the dispersion of proteinosomes can lead to changes in the membrane as it responds to a change in the osmolarity. These changes could be more easily accommodated for by the more porous and flexible membrane of the native proteinosome. In comparison, the change in the osmolarity led to the collapse and possible disassembly of the cationized proteinosome. Upon dilution of PURExpress to 0.33x, it was possible to observe intact BSA-NH_2_ proteinosomes which supported mRNA production by fluorescence microscopy. Analysis of these microscopy images showed that mRNA production was lower in the cationized proteinosome compared to the native proteinosomes (**Figure 2G, 2H, 2I**). Indeed, fluorescence spectroscopy experiments showed that expression within the BSA-NH_2_ proteinosome was reduced compared to the nat-BSA proteinosomes (**Figure 2, Figure S10**).

To determine, if the reduction of protein expression within the cationized proteinosome was due to the protein-PNIPAAm conjugate, we incubated 1mg.mL^-1^ of protein-PNIPAAm conjugate (nat-BSA-PNIPAAm or BSA-NH_2_-PNIPAAm) in non-diluted PURExpress with plasmid DNA coding for mRNA-Broccoli and mCherry. Microplate experiments showed no significant difference in the production of mRNA and protein in the presence of BSA-NH_2_-PNIPAAm compared to the nat-BSA-PNIPAAm (**Figure 3A**). Furthermore, comparisons to the control experiments without protein-PNIPAAm conjugate showed expression levels in the same order of magnitude. This suggests that protein expression was affected by the compartmentalization rather than by the individual protein-PNIPAAm conjugates. Given that both native and cationized proteinosomes used the same crosslinker the reduction in protein expression could be due to the ability for the expression machinery to diffuse across the membrane. To test this, we characterized the ability of FITC-labeled dextran (40 to 500 kDa) to permeate through the membranes of native and cationized proteinosomes (**Figure S11**). For native proteinosomes the molecule weight cut off for the proteinosomes prepared with BS(PEG)_2000_ was between 40 and 70 kDa for FITC-labeled dextran; for proteinosomes prepared with BS(PEG)_9_ the molecular weight cut off was less then 40 kDa. In comparison, the cationized proteinosomes showed low retention of 40 kDa FITC-labeled dextran. Our results show that the length of BS(PEG)_n_ affects the partitioning of molecules into the proteinosome as shown previously ^[23]^ and that protein cationization changes the permeability compared to the native protein. In this case cationized proteinosomes have a lower molecular weight cut off compared to the native proteinosomes. Even though the T7 polymerase is 99 kDa, our results showed mRNA production in both cationized and native proteinosomes which suggests that the polymerase can diffuse to the interior of the proteinosome (**Figure 2H**).

**Figure 3:**
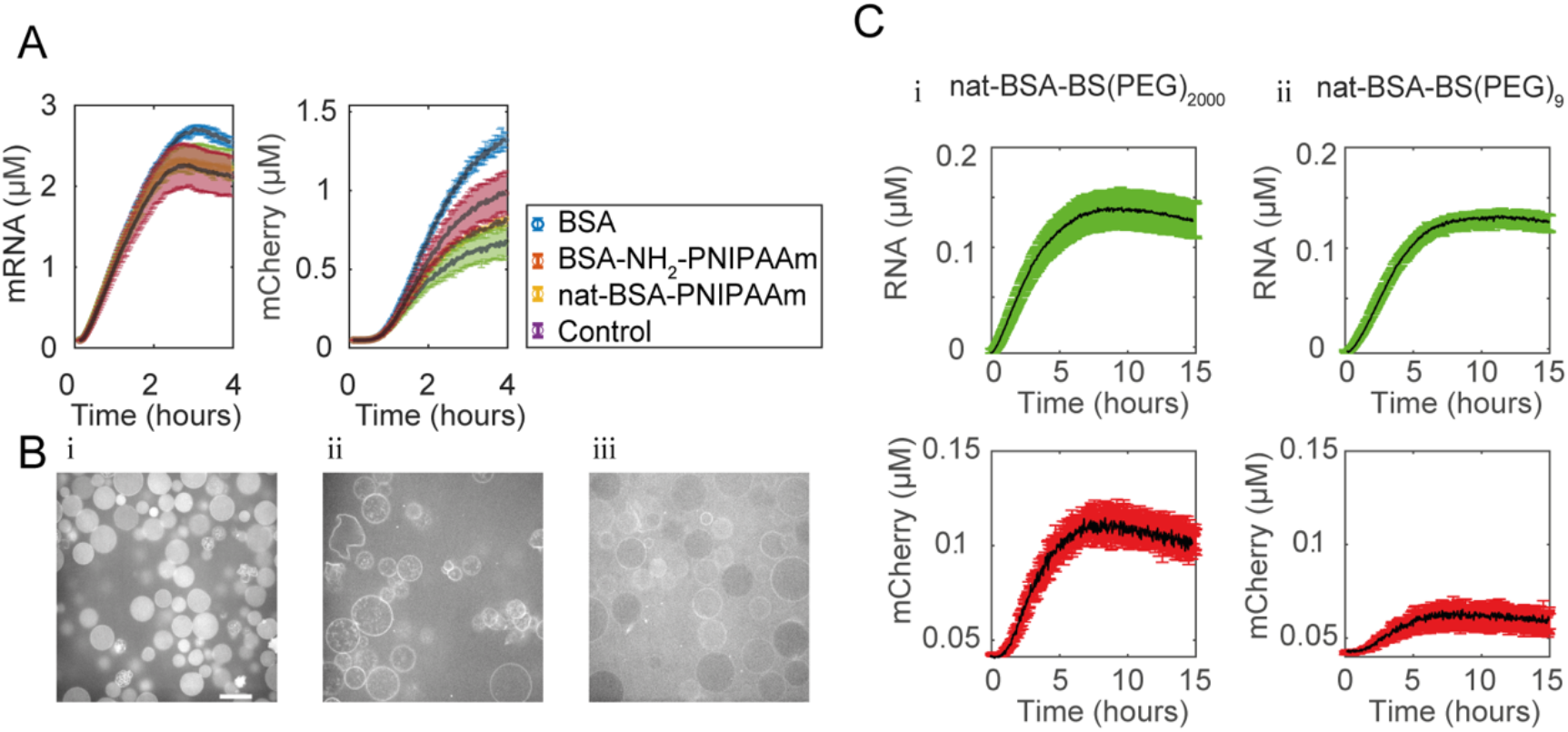
A) Cell free transcription (i) and translation (ii) in the presence of 1 mg.mL^-1^ of BSA (blue), BSA-NH_2_-PNIPAAm (orange) or nat-BSA-PNIPAAm (yellow) and a buffer control (purple) with PURExpress containing plasmid DNA (2.7 nM). Transcription (i) and translation (ii) were monitored via mRNA/DHFBI and mCherry respectively on a microplate reader. B) Permeability of labeled ribosomes into proteinosomes with DNA plasmid. Ribosomes were prepared in ribosome resuspension buffer and incubated with nat-BSA proteinosome crosslinked with BS(PEG)_2000_ (i); BSA-NH_2_ proteinosome crosslinked with BS(PEG)_2000_ (ii), and BSA-NH_2_ proteinosome crosslinked with BS(PEG)_9_ (iii). Cy5 labeled ribosome was imaged on a laser scanning confocal microscope with a 63x/1.3 NA Plan-Apochromat objective. C) Kinetics of transcription (green) and translation (red) in nat-BSA proteinosome crosslinked with BS(PEG)_2000_ (i) and BS(PEG)_9_ (ii) incubated with in PURExpress (0.5x) monitored on a microplate reader at 37°C. 3 repeats were performed and the shade were plotted as mean (black line) ±standard deviation. All microscopy images are representations of the results obtained from replicate experiments.

Furthermore, we tested the partitioning of fluorescently labeled ribosomes (**Figure S12**) into cationized and native proteinosomes (see materials and methods). Confocal microscopy images showed fluorescence from ribosomes within nat-BSA proteinosomes. In comparison ribosomes were aggregated within the interior of the BSA-NH_2_ proteinosomes or interacting at the surface (**Figure 3B**). This suggests that the ability for the ribosome to diffuse into the proteinosome where the DNA is located is affected by the cationization of the protein-PNIPAAm conjugate. This could be due to a decrease in pore size or surface interactions between the ribosomes and the proteinosomes. Our results suggest that by tuning the pore size of the proteinosome, protein expression levels can be regulated.

As it had previously been shown that changing the molecular weight of the BS(PEG)_n_ crosslinker could change the molecular weight cut off,^[23]^ we prepared native proteinosomes with a BS(PEG)_9_ crosslinker that encapsulated plasmid DNA (**Figure S13**). These nat-BSA-BS(PEG)_9_ proteinosomes were incubated with 0.5x PURExpress and transcription and translation were monitored on the microplate reader for mRNA and protein. Comparison between nat-BSA proteinosomes prepared with BS(PEG)_9_ and BS(PEG)_2000_ showed that the mRNA production was comparable but mCherry production was significantly reduced in the proteinosomes prepared with the smaller BS(PEG)_9_ (**Figure 3C**) when the PURExpress had been diluted by a half. Confocal microscopy images show that after 1 hr of incubation of the proteinosome with fluorescently labeled ribosomes showed a proportion of proteinosomes had excluded the ribosomes (**Figure 3B**). Together, our results showed that the exclusion or inclusion of the ribosome in the proteinosome regulates the amount of protein that is produced and the exclusion of ribosomes can be attenuated by crosslinker length and pore size of the proteinosomes or by molecular interactions with cationized protein-PNIPAAm conjugate.

It is interesting to note that nat-BSA proteinosomes prepared with DNA were stable in water for at least a year (**Figure S14**), at a range of pHs (**Figure S15**) and more stable than cationized proteinosomes at high osmolarity (**Figure S16**) which suggests that their applications can be widened beyond cell free transcription and translation as demonstrated here.

Therefore, to validate the versatility of the new method, we tested whether we could generate stable proteinosomes from GOx without any cationization. To this end, we conjugated PNIPAAm directly to commercially available GOx and crosslinked them with BS(PEG)_9_ as described for BSA. Imaging using optical microscopy and electron microscopy (**Figure S17**) showed stable proteinosomes. Further to this, we tested the reactivity of the nat-GOx proteinosomes by adding a dispersion of the proteinosomes to a phosphate solution containing Amplex Red and horseradish peroxidase (see material and methods). D-glucose was added to the reaction mixture at 0-50 µM concentrations and was immediately imaged on a microplate reader. Our results showed that nat-GOx proteinosomes remained active and provides a validation that this methodology can be readily transferred to other enzymes whilst maintaining enzymatic activity (**Figure S17**).

## Conclusions

In brief, we have adapted the methodology for the production of proteinosomes to integrate for the first time, cell free transcription and translation into crosslinked aqueous proteinosomes. We present a new method for the preparation of the proteinosome that directly conjugates the PNIPAAm to native BSA and glucose oxidase removing the chemically aggressive protein cationization step. Not only does this facile method provide a robust protocol which can be readily transferred between laboratories, it also reduces the number of positive charges on the protein polymer conjugate. We show that the reduction of the positive charges leads to native proteinosomes that are more stable to high salt environments compared to the cationized proteinosomes. This is significant because it means that cell free transcription and translation which requires high salt can be supported in these stabilized proteinosomes. Indeed, we show that transcription and translation is more efficient in native proteinosomes compared to cationized proteinosomes. Furthermore, decreased positive charge increases the diffusion of ribosome into the center of the proteinosome which is required for the translation process. In the case of the native proteinosome, the diffusion of ribosome into the interior of the compartment can be tuned by altering the pore size of the proteinosome by the length of the BS(PEG)n crosslinker. This provides a handle to tune the activity of the proteinsome by changing the one chemical in the methodology and without additional methodological steps.

The incorporation of cell free transcription and translation into proteinosomes provides an important tool for establishing synthetic cells with different architectural features. The porous membrane of the proteinosome and its ability to support transcription and translation could provide a minimal chemical model for isolating mRNA and DNA whilst allowing the diffusion of protein out of the proteinosomes and allowing the diffusion of resources into the proteinosome. The open nature of the proteinosome provides unique properties, *i*.*e*. resources can be continually supplied to the reaction to lengthen the time of the reaction. This is important for minimal systems which do not have the chemical complexity to autonomously produce energy from external resources that are required for the long-term sustenance of living systems.

Towards the goal of establishing synthetic cells from scratch the generation of multi compartments with different physical features can provide a route to replicate biological cellular architecture and to realise differential regulation of biochemical reactions. To this end, We show that the native proteinosomes can support higher order architectural structures with a “Russian doll” effect where we generate a 3-tiered system with a condensate encapsulated within a proteinosome (Figure S18) which is encapsulated within a liposome. from the inclusion of small compartments based on protein (**Figure S18**) (**Figure 4, Figure S19**). Additional design elements that include the transport of mRNA out of the proteinosome to separate transcription and translation could be an intriguing property to include into the proteinosome – that more closely resembles some of the features of the cell nucleus for example.

Integration of cell free transcription and translation in robust synthetic cellular chassis, as we have demonstrated, is critical not only in providing tangible systems to build synthetic cells but is a fundamental requirement to address a key challenge within synthetic cellular research. The challenge to extend synthetic cellular systems from basic research to applications in the real world. For this, facile and reproducible methods and the ability to control and tune enzymatic activities are a fundamental prerequisite.

## Supporting information

supplementary infomration

## Acknowledgements

T.Y.D.T. and M.G. acknowledge financial support from the MPG and MaxSynBio Consortium, jointly funded by the Federal Ministry of Education and Research (Germany) and the Max Planck Society and the MPI-CBG. We thank the Deutsche Forschungsgemeinschaft (DFG, German Research Foundation) under Germany’
ss Excellence Strategy – EXC-2068 – 390729961– Cluster of Excellence Physics of Life of TU Dresden and EXC-1056-Center for Advancing Electronics Dresden; Volkswagen Stiftung for generous funding. We acknowledge the Light Microscopy Facility, the Electron Microscopy facility and the Protein Expression and Purification (PEPC) at MPI-CBG for assistance. We also acknowledge Delong instruments for the assistance of the TEM image acquisition. We thank David T. Gonzales for the help in preparing liposome. We also acknowledge Archishman Ghosh for the purified PRM5. We thank the Center for Macromolecular Structure Analysis in the Leibniz Institute of Polymer Research, Dresden for GPC analysis of the PNIPAAm chain.

## Author contributions

M.G. and T.Y.D.T. conceived the research. M.G., D.W., M.W.-B., L.W. and J.S. contributed to the design of and the undertaking of the experiments. M.G., D.W., L.W. and J.S. analyzed the data. All authors contributed to the writing of the manuscript.

