## supplementary infomration for "Cell free expression in proteinosomes prepared from native protein-PNIPAAm conjugates"

**Materials:**

| Full Name | Abbreviation | Lot Nr. | Company |
|---|---|---|---|
| Dithiothreitol | DTT | R0862 | ThermoFisher scientific, USA |
| Bovine Albumin Serum | BSA | 421501j | Avantor, Germany |
| Evagreen fluorescence dna stain | EvaGreen | pcr379 | Jena Bioscience, Germany |
| Trizma base | Tris-HCl | T1503 | Sigma Aldrich Chemie GmbH, Germany |
| Potassium Chloride | KCl | 1.04936.1000 | Merck, Germany |
| Sodium Chloride | NaCl | 1.06404.1000 | Merck, Germany |
| Magnesium Chloride Hexahydrate | $MgCl_2$ | 1.05833.1000 | Merck, Germany |
| Ammonium Sulfate | $(NH_4)_2SO_4$ | 1.01217.1000 | Merck, Germany |
| Potassium Hydroxide | KOH | 1310-58-3 | Merck, Germany |
| PURExpress® Δ Ribosome Kit | | E3313 | New England Biolabs GmbH, UK |
| Magnesium Sulfate | $MgSO_4$ | M3409 | Sigma Aldrich Chemie GmbH, Germany |

| | | | |
|---|---|---|---|
| Magnesium acetate tetrahydrate | $Mg(OAc)_2$ | Css-210 | Jena Bioscience, Germany |
| 2-Mercaptoethanol | BME | M6250 | Sigma Aldrich Chemie GmbH, Germany |
| N-2-hydroxyethylpiperazine-N'-2-ethane sulphonic acid | HEPES | 9105.3 | Carl Roth, Germany |
| Potassium Phosphate monobasic | $KH_2PO_4$ | P8709 | Sigma Aldrich Chemie GmbH, Germany |
| Potassium Phosphate dibasic | $K_2HPO_4$ | P8584 | Sigma Aldrich Chemie GmbH, Germany |
| Sodium Hydrogen Carbonate | $NaHCO_3$ | 1.06329.1000 | Merck, Germany |
| Nuclease-free water | water | AM9937 | ThermoFisher scientific, USA |
| Glucose oxidase from Aspergillus niger, Type VII | GOx | G2133 | Sigma Aldrich Chemie GmbH, Germany |
| O,O′-Bis-[2-(N-succinimidyl-succinylamino)-ethyl]-polyethylenglykol | $BS(PEG)_{2000}$ | 713783 | Sigma Aldrich Chemie GmbH, Germany |
| BS(PEG)9 | $BS(PEG)_9$ | 21582 | Life Technologies GmbH, Germany |
| 2-ethyl-1-hexanol | oil | E0122 | Tokyo Chemical Industry UK Ltd., UK |
| Dimethyl sulfoxide | DMSO | D12345 | ThermoFisher scientific, USA |
| Peroxidase, horseradish | HRP | J60026 | ThermoFisher scientific, USA |
| Amplex Red reagent | Amplex Red | A12222 | Life Technologies GmbH, Germany |
| D-Glucose | glucose | G8270 | Sigma Aldrich Chemie GmbH, Germany |

| | | | |
|---|---|---|---|
| D-Sucrose | sucrose | 141621.1211 | AppliChem GmbH |
| DFHBI | DFHBI | SML1627 | Sigma Aldrich Chemie GmbH, Germany |
| 50% glutraldehyde for electron microscopy | GA | 002311601 | Serva, Germany |
| ChromoTek RFP-Trap Agarose | Bead | rta | Chromotek, Germany |
| Cyanine5-N-Hydroxysuccinimide-ester | Cy5-NHS | fab146454 | abcam, UK |
| *E-coli* Ribosome | ribosome | p0763s | New England Biolabs GmbH, UK |
| PURExpress in vitro protein syntehsis kit | PURExpress | E6800L | New England Biolabs GmbH, UK |
| slot grid | grid | cat# G2010-Cu | Science Services, UK |
| 400 mesh hexagonal grid | grid | G400H-Cu | Science Services, UK |
| Formvar 15/95 resin (polyvinyl Formal Resin) | | cat# 15800 | Science Services, UK |
| Uranyl Acetate | | Cat# 22400 | Electron Microscopy Sciences, Switzerland |
| Osmium Tetroxide | | Cat# 19190 | Electron Microscopy Sciences, Switzerland |
| Potassium hexacyanoferrate(II) trihydrate | | P9387-100G | Sigma Aldrich Chemie GmbH, Germany |
| EMbed-812 Resin | epon | cat# 14900 | Electron Microscopy Sciences, Switzerland |
| Lead Citrate | | Car# 17800 | Electron Microscopy Sciences, Switzerland |
| 4,4′-azobis (4-cyanovaleric acid) | ACVA | 2638-94-0 | Sigma Aldrich Chemie GmbH, Germany |
| 2-thiazoline-2-thiol | | 96-53-7 | Sigma Aldrich Chemie GmbH, Germany |

| | | | |
|---|---|---|---|
| 1,4-dioxane | | 123-91-1 | Sigma Aldrich Chemie GmbH, Germany |
| N, N′-dicyclohexylcarbodiimide | DCC | 538-75-0 | Sigma Aldrich Chemie GmbH, Germany |
| 4-(dimethylamino) pyridine | DMAP | 1122-58-3 | Sigma Aldrich Chemie GmbH, Germany |
| Diethyl Ether | | 60-29-7 | Sigma Aldrich Chemie GmbH, Germany |
| Sodium hydride, 60 % dispersion in mineral oil | NaH | 7646-69-7 | Sigma Aldrich Chemie GmbH, Germany |
| Ethyl acetate | | 141-78-6 | Fisher Chemical, USA |
| Propanethiol | | 107-03-9 | Sigma Aldrich Chemie GmbH, Germany |
| Carbon disulfide | $CS_2$ | 75-15-0 | Sigma Aldrich Chemie GmbH, Germany |
| Iodine | $I_2$ | 7553-56-2 | Sigma Aldrich Chemie GmbH, Germany |
| Sodium thiosulfate | $Na_2S_2O_3$ | 7772-98-7 | Sigma Aldrich Chemie GmbH, Germany |
| Sodium sulfate | $Na_2SO_4$ | 7757-82-6 | Sigma Aldrich Chemie GmbH, Germany |
| N-hexane | | 110-54-3 | Fisher Chemical, USA |
| N-isopropylacrylamide | NIPAM | 2210-25-5 | Sigma Aldrich Chemie GmbH, Germany |
| 2,2′-Azobis(2-methylpropionitrile) | AIBN | 78-67-1 | Sigma Aldrich Chemie GmbH, Germany |
| N, N′-Dimethylacetamide | DMAc | 127-19-5 | VWR Chemicals, Germany |
| Lithium chloride | LiCl | 7447-41-8 | Honeywell, USA |
| Poly(2-vinylpyridin) | PVA | 9017-40-7 | Polymer Standards Service, Germany |
| Chloroform-d | $CDCl_3$ | 865-49-6 | Eurisotop, USA |

| | | | |
|---|---|---|---|
| NMR tubes | | EP55.1 | Carl Roth Gmbh & Co. Kg, Germany |
| Sealing caps for NMR tubes | | HX59.1 | Carl Roth Gmbh & Co. Kg, Germany |
| L-Alanine | | A7627 | Sigma Aldrich Chemie GmbH, Germany |
| L-Arginine | | A5006 | Sigma Aldrich Chemie GmbH, Germany |
| L-Asparagine | | A0884 | Sigma Aldrich Chemie GmbH, Germany |
| L-Aspartic acid | | A9256 | Sigma Aldrich Chemie GmbH, Germany |
| L-Cysteine | | W326305 | Sigma Aldrich Chemie GmbH, Germany |
| L-Glutamic acid | | G1251 | Sigma Aldrich Chemie GmbH, Germany |
| L-Glutamine | | G3126 | Sigma Aldrich Chemie GmbH, Germany |
| Glycine | | G7126 | Sigma Aldrich Chemie GmbH, Germany |
| L-Histidine | | H8000 | Sigma Aldrich Chemie GmbH, Germany |
| L-Isoleucine | | I2752 | Sigma Aldrich Chemie GmbH, Germany |
| L-Leucine | | L8000 | Sigma Aldrich Chemie GmbH, Germany |
| L-Lysine | | L5501 | Sigma Aldrich Chemie GmbH, Germany |
| L-Methione | | M9625 | Sigma Aldrich Chemie GmbH, Germany |
| L-Phenylalanine | | P2126 | Sigma Aldrich Chemie GmbH, Germany |

| | | | |
|---|---|---|---|
| L-Proline | | P0380 | Sigma Aldrich Chemie GmbH, Germany |
| L-Serine | | S4500 | Sigma Aldrich Chemie GmbH, Germany |
| L-Threonine | | T8625 | Sigma Aldrich Chemie GmbH, Germany |
| L-Tryptophan | | T0254 | Sigma Aldrich Chemie GmbH, Germany |
| L-Tyrosine | | T3754 | Sigma Aldrich Chemie GmbH, Germany |
| L-Valine | | V0500 | Sigma Aldrich Chemie GmbH, Germany |
| Adenosine 5′-triphosphate disodium salt hydrate | ATP | A26209 | Sigma Aldrich Chemie GmbH, Germany |
| Cytidine 5′-triphosphate disodium salt hydrate | CTP | 30320 | Sigma Aldrich Chemie GmbH, Germany |
| Guanosine 5′-triphosphate sodium salt hydrate | GTP | 10106399001 | Roche, Switzerland |
| Uridine 5′-triphosphate trisodium salt dihydrate | UTP | 94370 | Sigma Aldrich Chemie GmbH, Germany |
| DiD (1,1'-Dioctadecyl-3,3,3',3'Tetramethylindodicarbocyanine, 4Chlorobenzenesulfonate Salt) | DiD | D7757 | Sigma Aldrich Chemie GmbH, Germany |
| 1-octanol | | 297887 | Sigma Aldrich Chemie GmbH, Germany |
| Egg PC (L-α-phosphatidylcholine) | Egg PC | 840051 | Avanti, Germany |
| L-Glutamic acid potassium salt monohydrat | K-Glutamate | G1149 | Sigma Aldrich Chemie GmbH, Germany |
| L-Glutamic acid hemimagnesium salt | Mg-Glutamate | 49605 | Sigma Aldrich Chemie GmbH, Germany |

| | | | |
|---|---|---|---|
| Folinic acid calcium salt hydrate | | F7878 | Sigma Aldrich Chemie GmbH, Germany, |
| pEXP5-NT/6xHis mCherry-F30-2xdBroccoli | Plasmid | 169233 | Addgene/home-made, USA |

**Preparation of protein-PNIPAAm conjugate**

PNIPAAm with a mercaptothiazoline terminal group was synthesized *via* RAFT polymerization as described previously[1–3]. PNIPAAm was characterized by Size Exclusion Chromatography (SEC) and Nuclear Magnetic Resonance Spectroscopy (NMR) as previously described[4]. The dispersity ($Đ = M_w/M_n$; $M_n$ is the number average molecular weight; $M_w$ is the weight average molecular weight of block copolymers and were detected using a size exclusion column equipped with a MALS detector (MiniDAWN-LS detector, Wyatt Technology, USA) and a viscosity/refractive index (RI) detector (ETA-2020, WGE Dr. Bures, Germany). The column (PL MIXED-C with a pore size of 5 µm, 300 × 7.5 mm) and the pump (HPLC pump, Agilent 1200 series) were from Agilent Technologies (USA). 2% vol water in DMAc and 3 $g.L^{-1}$ of LiCl were flowed at a rate of 0.5 $mL.min^{-1}$ to elute the polymer. PVA was used as a standard at 2 $mg.mL^{-1}$ after filtration through a 0.2 µm filter. The data were processed using Cirrus GPC offline GPC/SEC software (version 2.0). The synthesized PNIPAAm chain was then measured with a number average molecular weight ($M_n$) of 14000 $g.mol^{-1}$ and a weight average molecular weight of block copolymer ($M_w$) of 17000 $g.mol^{-1}$ resulting in a dispersity index ($Đ = M_w/M_n$) of 1.12 unless otherwise stated.

Cationized protein-PNIPAAm conjugates (BSA-$NH_2$-PNIPAAm and GOx-$NH_2$-PNIPAAm) were prepared as previously described[1,3]. Briefly, native protein (BSA or GOx was cationized, *via* carbodiimide chemistry, with aminohexane to increase the number of available amine groups. The cationized protein (BSA-$NH_2$ or GOx-$NH_2$) was reacted with mercaptothiazoline-activated PNIPAAm at a molar ratio of 1:8.3 for BSA-$NH_2$:PNIPAAm and 1:19 for GOx-$NH_2$:PNIPAAm in 0.2 M $NaHCO_3$. Native protein-PNIPAAm conjugates (nat-BSA-PNIPAAm and nat-GOx-PNIPAAm) were prepared by directly mixing native protein solution (1 $mg.mL^{-1}$) with PNIPAAm in 0.2 M $NaHCO_3$ solution at a molar ratio of 1:8.3 (BSA: PNIPAAm) or 1:11.7 (GOx:PNIPAAm).

In all instances, the solution of protein and polymer was incubated with shaking at 4˚C for 16 hours, then filtered with 0.22 µm PVDF filter into a 25 kDa dialysis tubing (POR, Thermofisher, USA). The sample was then dialyzed against 0.2x PBS at 4 ˚C, the buffer was refreshed every hour for 3 hours to remove any unconjugated polymer. After dialysis, the native and cationized protein conjugate was concentrated to a tenth of the original volume, aliquoted, snap frozen in liquid nitrogen and stored at -20 ˚C until further use.

**Characterization of the protein-PNIPAAm conjugate:**

**Confirmation of protein-PNIPAAm conjugation**

**Dynamic light scattering (DLS)**

The size of native BSA and nat-BSA-PNIPAAm were measured on a Zetasizer Nano ZSP (Malvern Panalytical, UK). For DLS measurements native BSA, BSA-PNIPAAm and BSA-$NH_2$ at 1 mg.mL$^{-1}$ were loaded into a ZEN0040 plastic micro cuvette and measured at 20˚C and 37˚C. 3 technical repeats were performed with 3 measurements per round.

**Laser induced liquid beam ion desorption mass spectrometry (LILBID-MS[5])**

Solutions of BSA-$NH_2$-PNIPAAm and nat-BSA-PNIPAAm was dispensed *via* a piezo-driven droplet dispenser head (MD-K-130 from Microdrop Technologies GmbH, Germany) to generate sample droplets of about 50 µm diameter with a repetition rate of 10 Hz. These microdroplets were transferred to high vacuum and irradiated by a mid-IR laser pulse. The laser employed was a Nd:YAG laser whose wavelength could be tuned by a $LiNbO_3$ optical parametric oscillator to approximately 2.8 µm. The pulse length was 6 ns with a maximum energy of 23 mJ. The laser power was measured by an optical power meter (PM100D, Thorlabs, Germany). The laser excites the asymmetric O-H stretch vibration of water, which subsequently leads to a rapid expansion of the droplet. During this process solvated ions are released and accelerated into a Wiley-McLaren-type time-of-flight analyzer. By applying a voltage difference between the enclosing ion lenses (repeller and extractor), ions of the corresponding charge are led into the grounded flight tube and guided towards the detector *via* reflection.

The Daly-type detector is optimized for the detection of high m/z ions. The voltage of the repeller and extractor were set to -4 kV. The acceleration of the ions was initiated by a pulsing repeller at – 6.6 kV for 370 µs, 15 µs after droplet irradiation. Einzel lenses in front of and after the reflectron were set to − 3.0 kV, the reflection itself to − 7.2 kV. Post-acceleration was set to + 17 kV at the MCP impact surface. Spectra processing was undertaken with OriginPro 2021 by OriginLab Corporation.

**Determination of protein-polymer concentrations**

The concentration of native and cationized protein-PNIPAAm conjugate and the molar ratio of protein to polymer was determined by UV spectroscopy. To do this, calibration curves of PNIPAAm and native protein (BSA and GOx) were obtained by serial dilutions, in water, of known concentrations of polymer and protein. For PNIPAAm the concentration was determined by dry mass and absorbance at 300 nm of 50 µL of sample.

For the protein calibration, the Pierce bicinchoninic protein assay kit (Thermo Fisher Scientific, USA) was used as per the manufacturers' instructions with incubation of the assay with known concentrations of protein at 37 ˚C for 1 hour and measuring the absorbance at 561 nm for 100 µL of sample.

To determine the molar ratio and concentration of protein-PNIPAAm conjugate, a solution of conjugate was diluted 25 or 50 times to ensure that the sample was in a comparable range to the calibration curve. 50 µL was loaded into a 96 well plate and the absorbance measured at 300 nm to measure the PNIPAAm concentration. 50 µL of the BCA protein assay was then added to the sample, incubated at 37 ˚C for 1 hour. The absorbance measured at 561 nm was used to determine the protein concentration. The molar ratio of protein to polymer was then calculated.

All microplate reader experiments were conducted on Tecan Spark 20M multimode microplate reader (Tecan AG, Switzerland).

**Proteinosome preparation**

Proteinosomes were prepared from native or cationized protein-polymer conjugates. BSA-$NH_2$- PNIPAAm (70 µM), nat-BSA-PNIPAAm (74 µM), nat-GOx-PNIPAAm (50 µM**)** and GOx-$NH_2$-PNIPAAm (50 µM) were mixed on ice with or without plasmid DNA in 0.1 M $NaHCO_3$ to produce an aqueous solution of final volume of 60 µL. BS$(PEG)_{2000}$ (freshly prepared at 100 mM) or BS$(PEG)_9$ (100 mM) dissolved in anhydrous DMSO was mixed to a final concentration of 2 mM (BS$(PEG)_{2000}$) or 0.25 mM ( BS$(PEG)_9$) in 2-ethyl-1-hexanol (oil) at room temperature by vortexing. 1 mL of the oil mixture was transferred to the aqueous solution with a long-tip 1 mL pipette and pipetted 10 times at a speed of two times per second to generate an emulsion. After emulsification, an additional 1 mL of 2-ethyl-1-hexanol mixture was added and the mixture was stored at 8 ˚C overnight to allow complete crosslinking.

2-ethyl-1-hexanol was exchanged with water by firstly centrifuging the emulsion for 5 seconds in a bench centrifuge at room temperature then removing the supernatant by pipetting. 1 mL of cold (-20 ˚C) 75% ethanol was added and the pellet was fully resuspended by cautious pipetting. The solution was then transferred to a clean Eppendorf tube and left in the fridge for 2-3 hours. The sample was washed by repeated steps of centrifugation at 3000 rcf for 3 min at room temperature for BSA and at 8˚C for GOx based proteinosomes followed by removal of the supernatant and then resuspension of the proteinosome pellet with 1 mL of ice-cold 50%, 25% ethanol and 2x ultrapure water by careful pipetting. The proteinosomes were then stored in the fridge until further experiments.

The BSA-$NH_2$-PNIPAAm and GOx-$NH_2$-PNIPAAm proteinosomes were prepared as previously described[1], with the exception that the proteinosomes were washed and transferred to water by centrifugation rather than by dialysis.

**Cell free expression (CFE) within proteinosomes**

pEXP5-NT/6xHis mCherry-F30-2xdBrocoli was prepared as previously described[6].

PURExpress *in vitro* protein expression kit was prepared as described by the manufacturer's instructions with and without proteinosomes containing plasmid DNA. For plasmid DNA coding for the mRNA aptamer (Broccoli) the mRNA binding fluorophore DHFBI (10 µM) was added to the reaction mix. For experiments where the PURExpress was diluted, components A and B were diluted in ribosome buffer (20 mM HEPES-KOH pH 7.6, 30 mM KCl, 10 mM $Mg(OAc)_2$) to a final ratio of 1:2 (0.5x) or 1: 3 (0.33x) (PURExpress: Buffer). 4 µL of sample was loaded in a 1536 multi-well plate (flat black, Greiner, Germany), proteinosomes were added to the PURExpress mixture and the plate was centrifuged at 3000 rcf for 3 min at room temperature before loading into the microplate reader and measured at 37 ˚C. Transcription and translation were monitored by fluorescence spectroscopy with $\lambda_{exc}$ = 470/20 nm $\lambda_{emiss}$ = 532/20 nm to detect DHFBI (mRNA) and $\lambda_{exc}$ = 585/10 nm $\lambda_{emiss}$ = 610/10 nm to detect reporter protein mCherry. For experiments containing the RFP trap beads, the beads were diluted 10 times in water and then loaded into the CFE mixture to a final dilution of 1:100.

The plate was imaged on a confocal microscope at the end of the experiment, after an overnight reaction. Proteinosomes were labeled with Cy5-NHS with a final concentration of 0.01 $mg.mL^{-1}$.

**Note on preparation of proteinosomes for cell free expression:** DNA plasmid was loaded into the proteinosomes during the preparation step. After transfer to water cell free expression system was added to the proteinosome dispersion to initiate gene expression. There were a number of reasons to design the experiment in this way compared to including the cell free expression system during the proteinosome preparation. Firstly, this avoids any potential issues from chemical crosslinking of the proteins from the expression system during the crosslinking step, secondly, it is likely that the cell free expression system would be rendered inactive during the ethanol wash step required to transfer the proteinosomes into water. Furthermore, key expression machinery would be lost through the membrane during the water transfer step due to the permeability of the membrane. Finally, addition of the cell free expression system during the preparation step will trigger gene expression, therefore the addition of the expression system into preformed DNA proteinosomes provides the control over the initiation of gene expression.

**Optical imaging of proteinosomes**

Multi-well plates containing proteinosomes and cell free expression system were imaged at 37˚C with an Andor IX83 inverted microscope equipped with

Yokogawa CSU-W1 spinning disk (Yokogawa, Japan), an Andor iXon ultra 888 Monochrome EMCCD camera and Andor iQ3 (3.6.2, Andor, UK) for imaging acquisition. UPLFLN20x/0.5 NA objective (Zeiss, Germany) was used for the time lapse imaging together with a Z-drift compensation system. 488 nm laser and $\lambda_{exc}$ = 480/40 nm $\lambda_{emiss}$ = 525/50 nm, LP 514 nm was used for imaging EvaGreen or DHB1; 561 nm laser and $\lambda_{emiss}$ = 685/40 nm, LP 514 nm was used for imaging mCherry.

A UPLSAPO60xS2/1.3 NA silicon oil objective (Zeiss, Germany) was used to compare different proteinosome groups in different buffers (ribosome buffer at pH 2 and 7.6; 0.2 M $NaHCO_3$ at pH 8.4 and ribosome buffer containing 1 M sucrose at pH 7.6) to test the stability of the proteinosomes under different conditions. For these experiments imaging of the proteinosomes was undertaken with EvaGreen or DFHBI.

For kinetics experiments that contained red fluorescent protein (RFP) nanotrap beads, two different focal planes with an approximate distance of 40 µm distance in the Z direction were chosen for imaging.

**Image analysis**

Optical images were analyzed using FIJI[7] with the StackRecJ plugin. To analyze a time series, the stacks were first drift corrected using an ImageJ based plugin StackRecJ with 'Translation' mode. Then the proteinosome or agarose beads were located with DFHB1 or protein fluorescence using the last frame of the image series. After a series of processes including; Gaussian blurry, thresholding, binary and particle analysis, the size of the proteinosomes and fluorescence from all channels were extracted. The data was averaged and plot using Matlab.

**Electron microscopy imaging of proteinosome**

**Negative staining:**

Proteinosomes were diluted 10 times in ultrapure water and loaded onto a 400-mesh hexagonal grid with formvar film and a thin layer of carbon coating. The grid was left over a pre-wetted filter paper for 3 min. The grid was blotted with filter paper wedge and incubated with 1% uranyl acetate (Polyscience Europe GmBH)water solution for 30 seconds. The solution was then removed with the edge of a filter paper, then the grid was transferred to be imaged on a Tecnai 12 TEM (Philips/FEI/ Thermofisher Scientific, 100 kV, USA) with a digital F416 CMOS camera (TVIPS, Germany).

**Proteinosome resin embedding:**

Sedimented and concentrated proteinosomes were suspended in water and contrasted with 1% osmium tetroxide/1.5% potassium hexacyanoferrate (II) trihydrate (Sigma-Aldrich)for 30 min on ice. The mixture was washed with water 3 times, the aqueous solution was subsequently replaced by 50%, 70%, 80%, 90% and 96% ethanol *via* centrifugation at 2000 rcf for 5 min each. Between each wash step the samples were incubated for 15 min on ice. The proteinosomes were resuspended in 100% ethanol by 3 rounds of 1 min centrifugation at 2000 rcf and 30 min incubation at room temperature.

The dehydrated proteinosome were gradually infiltrated with 1:2 Epon-812 replacement (embed-812,Science Services, Germany) / ethanol and incubated overnight, 2:1 Epon/ethanol for 6 hours and then overnight again with pure Epon. Centrifugation at 6000 rpm (3427 rcf) for 10 min took place between each infiltration step. A small piece of the pellet was polymerized in a silicon mould at 60 ˚C for 48 hours and sectioned to 70 µm slices on a Leica UC7 ultramicrotome. The sections were post-stained with aqueous uranyl acetate and lead citrate prior to imaging using a Tecnai 12 (Philips/FEI/ Thermofisher Scientific, 100 kV, USA) with a digital F416 CMOS camera (TVIPS,Germany) at 100 kV.

Electron microscopy visualization of the native proteinosome sample was performed using an All-in-One electron microscope LVEM 25 E (Delong Instruments, Czech Republic) in a transmission (TEM) regime at an operating voltage of 25 kV and scanning transmission (STEM) regime without post-staining of the sections.

**Partitioning of dextran-FITC**

Dextran-FITC (40, 70, 150, 250, 500 kDa) was diluted to a final concentration of 0.1 $mg.mL^{-1}$ and mixed with proteinosomes in in water. The fluorescence intensity within the proteinosomes was extracted using FIJI and compared to the background fluorescence. Data from twenty proteinosomes were averaged in each experiment.

**Fluorescent labeling and characterization of ribosome**

Cy5-NHS was used to label ribosome as previously described[8]. In short, 3 molar excess Cy5-NHS was mixed with E coli ribosome and incubated on ice for 20 min, then 37 ˚C for 20 min. After an overnight incubation in the cold room, the mixture was layered with buffer (20 mM HEPES-KOH pH7.6, 30 mM KCl, 10 mM $Mg(OAc)_2$, 7 mM 2-mercaptoethanol and 2 M sucrose) and ultracentrifuged at 54000 rpm with rotor TLA 100.3 (157972 rcf). The pellet was washed briefly with the ribosome storage buffer (20 mM HEPES-KOH pH 7.6, 30 mM KCl, 10 mM $Mg(OAc)_2$, 7 mM 2-mercaptoethanol) and resuspended in the same storage buffer at twice its original volume. The labeled ribosome was aliquoted and stored at -20˚C before use.

To test the integrity of the ribosome after fluorescence labeling, mass photometry was performed on Refeyn TwoMP (Refeyn, UK). BSA, immunoglobulin G (IgG) and thyroglobulin were used to set the calibration curve. Ribosome and Cy5-labeled ribosome were diluted in ribosome buffer (20 mM HEPES-KOH pH 7.6, 30 mM

KCl, 10 mM $Mg(OAc)_2$) with a dilution factor of 13300x and 4000x respectively to achieve optimal counting events. Both samples were measured with 3 repeats.

**Partitioning of ribosomes.**

For the partitioning experiments, labeled ribosome was added to proteinosomes in ribosome diluting buffer or in PURExpress ΔRibosome Kit. All samples were measured using a spinning disk microscope with a UPLSAPO60xS2/1.3 NA silicon oil objective, 638 nm laser and $\lambda_{emiss}$ = 685/40 nm filter set at 37 ˚C.

**Glucose oxidase assay**

The enzymatic activities of nat-GOx and GOx-$NH_2$ proteinosomes were determined by fluorescence spectroscopy. Nat-GOx and GOx-$NH_2$ proteinosomes proteinosomes were obtained from an overnight sedimented sample, and diluted in water 15 times. The dispersion of proteinosomes was added to a solution of 50 µM Amplex Red, 0.1 unit.mL$^{-1}$ HRP in 50 mM potassium phosphate buffer (pH 7.4). D-glucose was added to the reaction mixture at 0- 50 µM concentrations and 100 µL was immediately loaded into a flat bottom 96 well plate (Greiner, Germany). The well plate was loaded into a microplate reader and resorufin production was measured with $\lambda_{exc}$ = 530/15 nm and $\lambda_{emiss}$ = 590/15 nm.

**Encapsulation experiments**

Proteinosome containing membrane free compartments were conducted using purified PRM5, which was a gift from Dr. A. Ghosh. Plasmid contained nat-BSA-BS$(PEG)_{2000}$ proteinosome was mixed with EvaGreen (10 µM final concentration) and PRM5 (12 µM final concentration) in Ribosome dilution buffer and imaged on microscopy.

All samples were imaged on a laser-scanning confocal microscope with Zeiss LSM 880 inverted single-point scanning confocal microscope equipped with a 32 GaAsP photomultiplier tube channel spectral detector and 40x objective (C-Apochromat 40× NA 1.2 W objective, Zeiss). 488 nm laser and $\lambda_{emiss}$ = 499-535 nm was used for imaging EvaGreen; 633 nm laser and $\lambda_{emiss}$ = 640-720 nm was used for imaging DiD.

Inverse emulsion phase transfer encapsulation was performed a modified protocol as previously described[2]. In brief, a lipid oil phase consisting of 0.3 mM Egg PC lipid and 1.5 µM DiD was prepared in mineral oil and mixed by 1 hour sonication at 37 ˚C in a water bath (Bandelin, Germany). 6.25 µL PURExpress was mixed with 1 µL non diluted mCherry plasmid and 0.1 mg.mL$^{-1}$ 2000 kDa Dextran-FITC contained nat-BSA-BS$(PEG)_{2000}$ proteinosome and added into a 1.5 mL Eppendorf tube with 200 µL lipid oil phase. The emulsification was generated by dragging the Eppendorf tube over an Eppendorf tube rack for 15 times. 100 µL of the emulsion was then gently layered on top of a 40 µL lipid oil phase solution and 100 µL outer feeding buffer solution (5 mM ATP, 1.5 mM CTP, 1.5 mM UTP, 3 mM GTP, 0.5 mM amino acids, 1.5 mM DTT, 0.02 mM folinic acid, 280 mM K-glutamate, 20 mM Mg-glutamate, 150 mM HEPES, 200 mM glucose and 1.5 mM Spermidine). The Eppendorf tube was then centrifuged at 4500 rcf for 20 min to produce liposomes. The samples were transferred to a 384 well plate.

PRM5 contained proteinosome was generated by adding final concentration of 10 µM PRM5 into the mCherry plasmid contained nat-BSA-BS$(PEG)_{2000}$ proteinosome in 1x ribosome storage dilution with 500 mM sucrose. 12.5 µL of the aqueous phase was emulsified as above mentioned and loaded to Eppendorf tube containing an oil/lipid mixture and an outer solution containing 500 mM glucose in 1x ribosome dilution buffer. Liposomes were prepared by centrifugation at 4500 rcf at 10 min.

Multi-well plates containing liposomes were imaged at 37˚C with an Andor IX83 inverted microscope equipped with Yokogawa CSU-W1 spinning disk (Yokogawa, Japan), an Andor iXon ultra 888 Monochrome EMCCD camera and Andor iQ3 (3.6.2, Andor, UK) for imaging acquisition. UPLSAPO60xS2/1.3 NA silicon oil objective (Zeiss, Germany) was used with 488 nm laser, $l_{exc}$ = 480/40 nm $l_{emiss}$ = 525/50 nm, LP 514 nm and 561 nm laser and $l_{emiss}$ = 685/40 nm, LP 514 nm.

**Supplementary Figures**

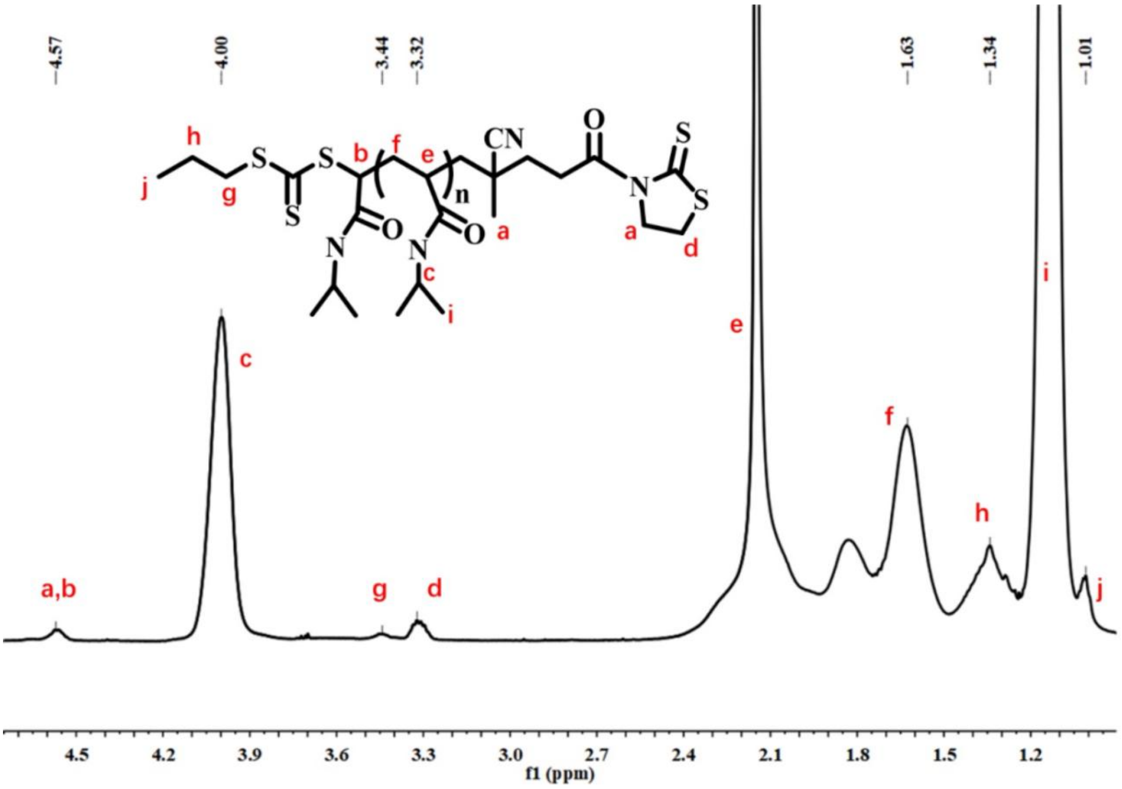


**Figure S1: $^1$H-NMR spectrum of mercaptothiazoline-activated PNIPAAm in $CDCl_3$.** The data was analyzed using MestReNova.

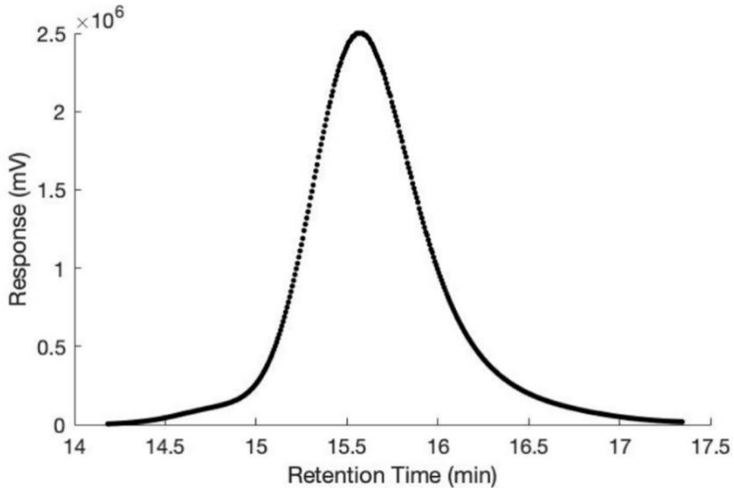


**Figure S2: Size exclusion chromatography (SEC) profile of PNIPAAm.** The PNIPAAm chain had a number average molecular weight ($M_n$) of 14000 g.mol$^{-1}$ and a weight average molecular weight ($M_w$) of 17000 g.mol$^{-1}$ resulting in a dispersity index (**Đ** = $M_w/M_n$) of 1.21. The molecular weight was a relative estimation.

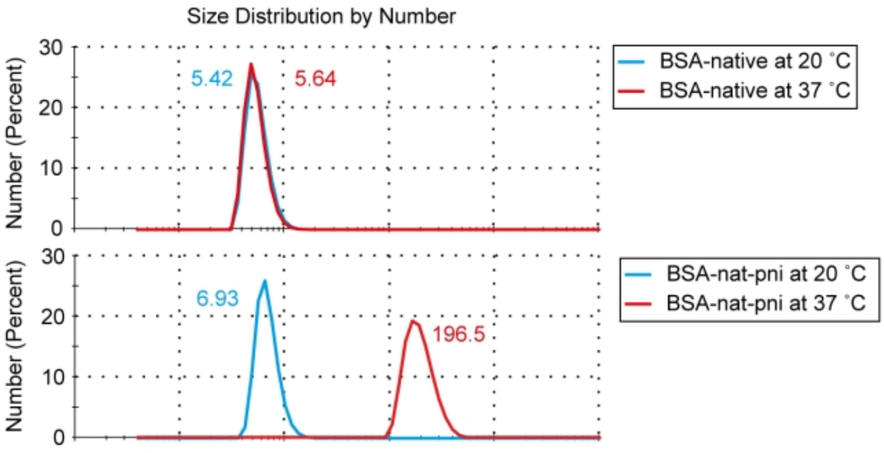


**Figure S3: DLS analysis of BSA with different modifications.** Native BSA, nat-BSA-PNIPAAm were prepared at 1 mg.mL$^{-1}$ and measured on a Zetasizer Nano ZSP at 20 ˚C and 37 ˚C. DLS show increase size of protein-polymer conjugate at 37 ˚C due to aggregation of conjugates driven by PNIPAAm collapse. 3 repeats were performed for each sample at each temperature.

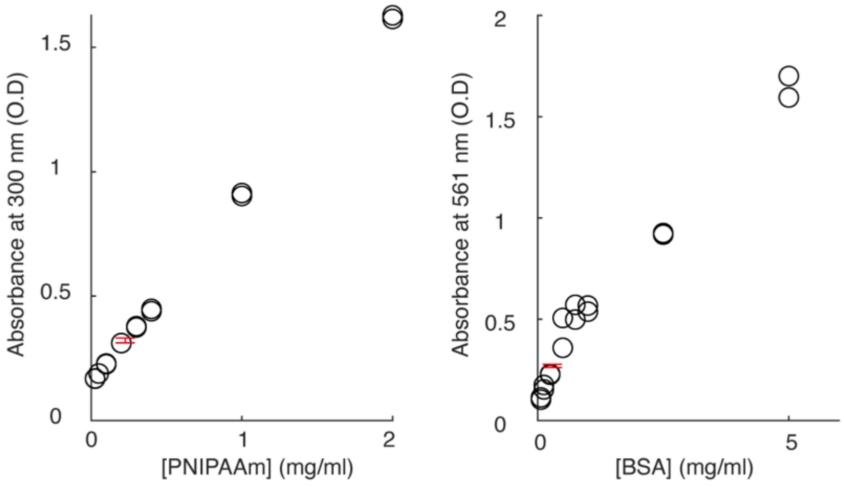


**Figure S4: Calibration curves used to determine concentration of nat-BSA-PNIPAAm.** Left: A serial dilution of PNIPAAm was used to calibrate the PNIPAAm concentration by UV absorbance at 300 nm; right: The same sample was stained with BCA reagent and measured with absorbance at 561 nm to determine the protein concentration. Tecan 20M plate reader was used for the measurement and 2 repeats were undertaken for the calibration (black open circles), 4 repeats for the sample (red error bars) were performed to obtain the mean concentration, points are the mean, error bars are standard deviation from the mean of the data.

**Table S1: Molar ratio of PNIPAAm with protein** determined by UV-Vis absorbance assays.

| **Molar ratio** | **GOx** | **GOx-NH$_2$** | **BSA** | **BSA-NH$_2$** |
|---|---|---|---|---|
| Initial PNIPAAm: Protein ratio | 11.7:1 | 19.0:1 | 8.3:1 | 8.3:1 |
| Calculated PNIPAAm:Protein ratio | 4.3:1 | 4.7:1 | 5.7:1 | 7.6:1 |

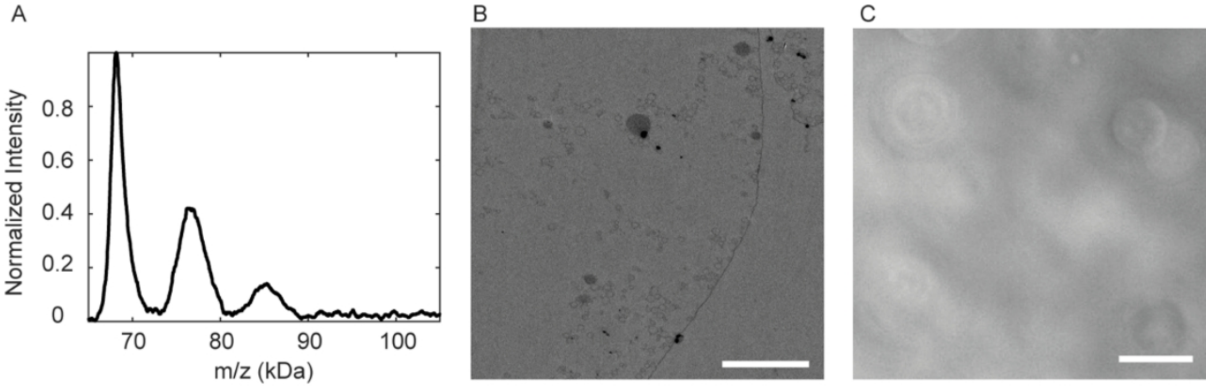


**Figure S5**: **Analysis of cationized conjugate and proteinosomes**. A) Mass-spectra of BSA-NH$_2$-PNIPAAm. B) EM image of cross-sections of embedded BSA-NH$_2$ proteinosome. Scale bar: 1 µm. C) BSA-NH$_2$ proteinosome imaged under phase contrast microscopy. Scale bar: 20 µm.

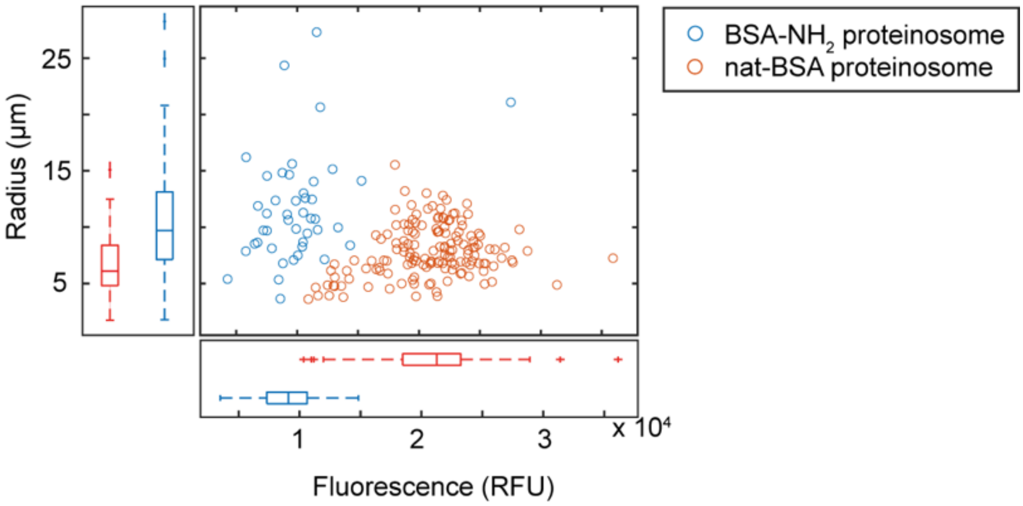


**Figure S6**: Nat-BSA proteinosomes have higher encapsulation efficiency and smaller proteinosomes compared to BSA-$NH_2$ proteinosomes. Confocal microscopy images of nat-BSA and BSA-$NH_2$ proteinosomes encapsulating plasmid DNA were compared using confocal microscopy. The fluorescence was from imaging in the EvaGreen channel for stained plasmid DNA. Two samples for each kind were imaged.

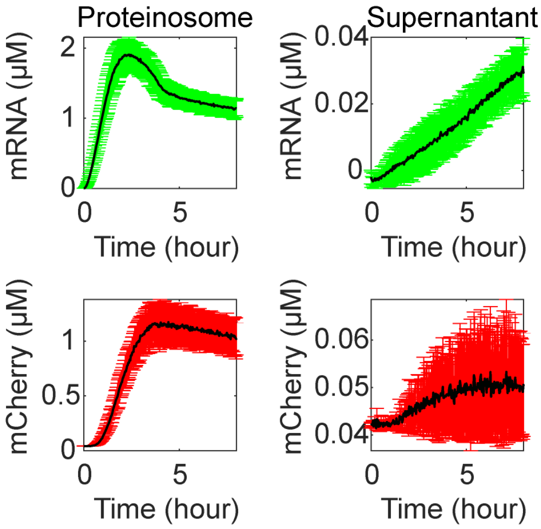


**Figure S7 Cell free expression with the supernatant from sedimented nat-BSA proteinosomes.** The proteinosomes with plasmid DNA were stored in water in an Eppendorf tube. After 1 month, the supernatant was removed and incubated with PUREpress at 37˚C to monitor the transcription *via* DHFBI and translation of mCherry using a Tecan 20M well plate reader (right). The proteinosome was added to PURExpress and the transcription and translation monitored *via* DHFBI and translation of mCherry using a Tecan 20M well plate reader (left). The results show that there was limited DNA leakage from the proteinosomes into the supernatant. 3 repeats were performed and the shade were plotted as mean (black line) ±standard deviation.

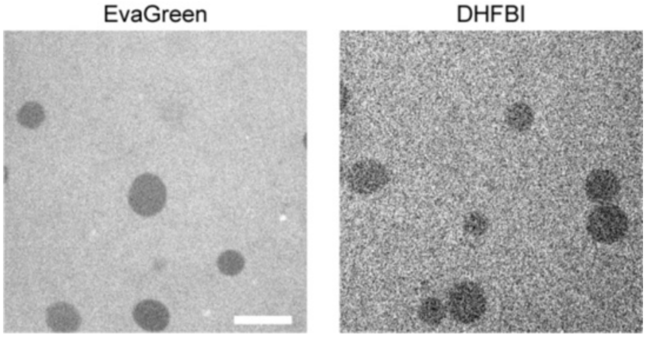


**Figure S8 Cell free expression reaction with empty proteinosomes.** 2.7 nM 2xBrocolli-mCherry plasmid was incubated with nat-BSA-BS(PEG)$_{2000}$ proteinosomes in PUREpress at 37˚C for 3 hours with either EvaGreen (left) or DHFBI (right). At the end of the reaction, the samples were imaged on a spinning disk confocal microscope using a 20x/0.5 NA objective at 37˚C. Both DNA (left) and mRNA (right) show low penetration into the proteinosomes. Scale bar 20 µm.

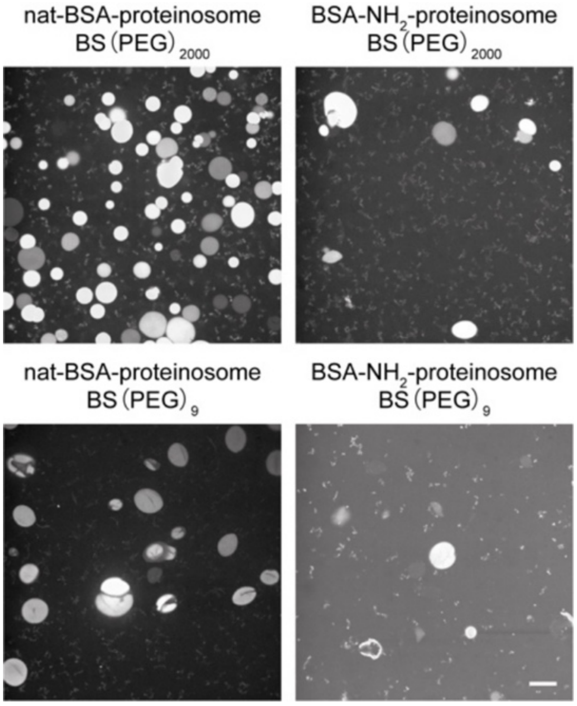


**Figure S9: Proteinosome morphology after cell free expression reaction in 1x PURExpress.** The proteinosomes encapsulating plasmids were stained with

EvaGreen dye and imaged after 3 hours in cell free expression reaction at 37˚C on a spinning disk confocal microscope. Scale bar 20 μm.

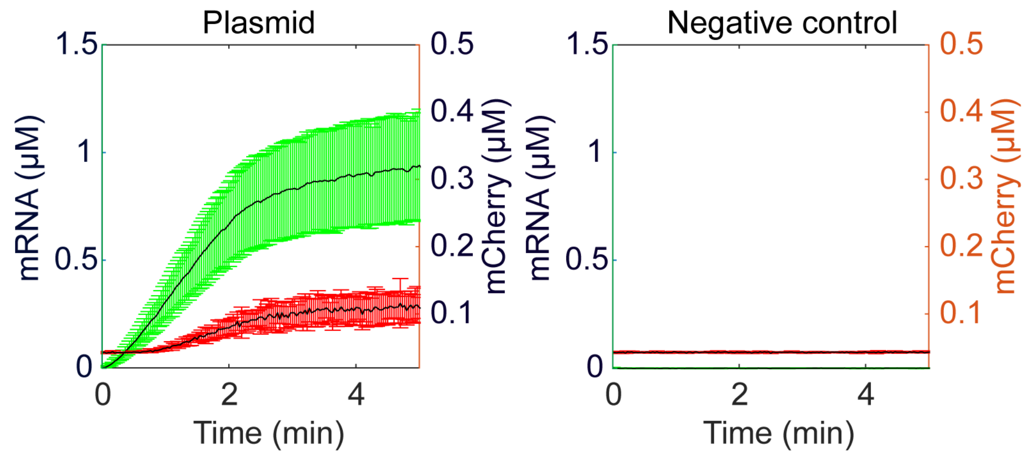


**Figure S10 Plasmid controls in diluted PURExpress mixture.** Plasmid (left: mCherry plasmid; right: negative control plasmid from NEB PURExpress kit) were incubated PURExpress. (0.33x). Fluorescence of mRNA/DHFBI and mCherry were monitored on a Tecan Spark 20M microplate reader. 3 repeats were performed and the shade were plotted as mean (black line) ±standard deviation.

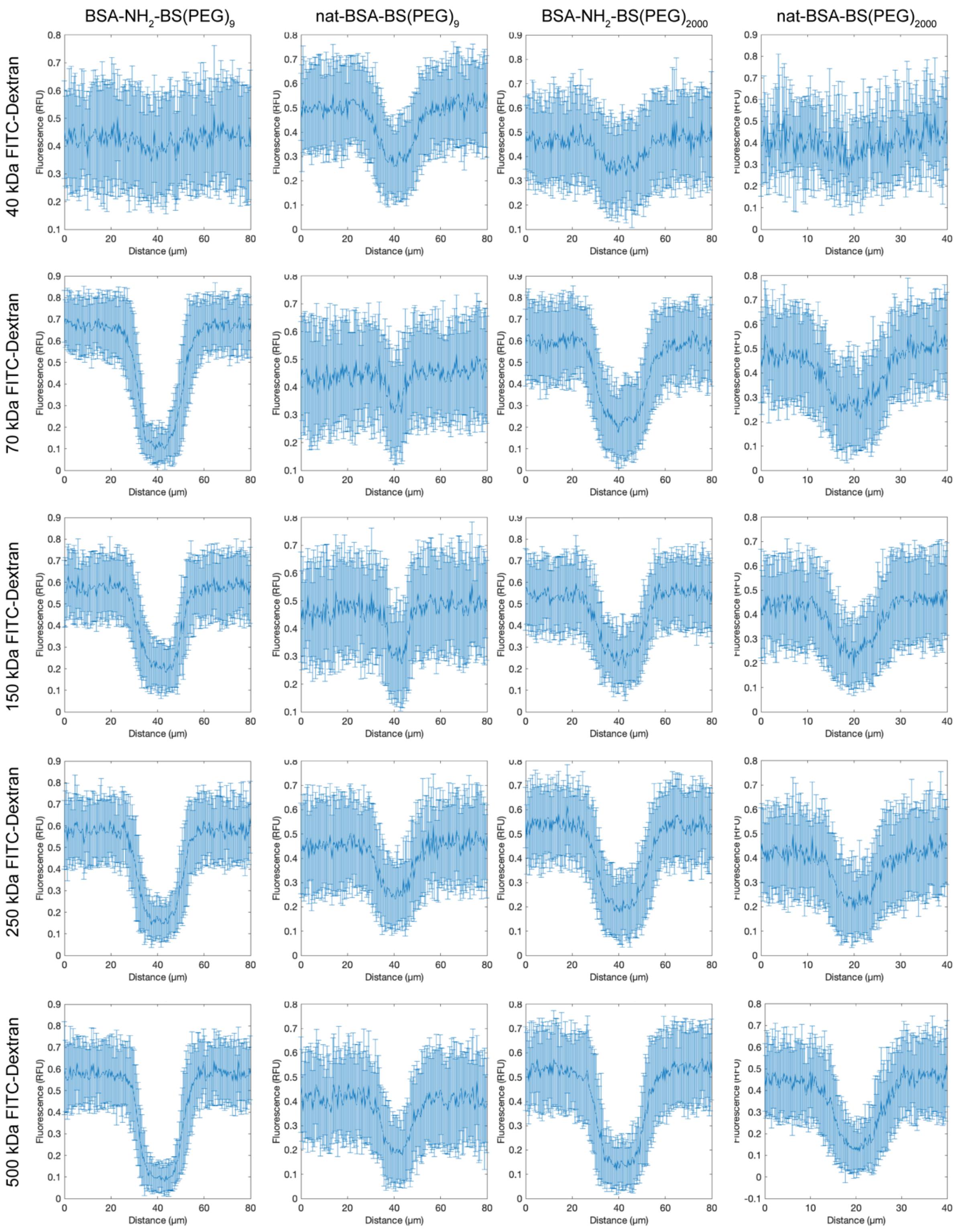


**Figure S11 Permeability of different proteinosomes to FITC-dextrans of varying molecular weight .** Dextran-FITC at different molecular weights were incubated with proteinosomes and imaged at 37˚C on a laser scanning confocal microscopy. Plots were obtained from line profiles from at least 20 proteinosomes from two independent experiments which were averaged and normalized for each sample, mean and standard deviations were plotted in the figures as solid line and error bars respectively.

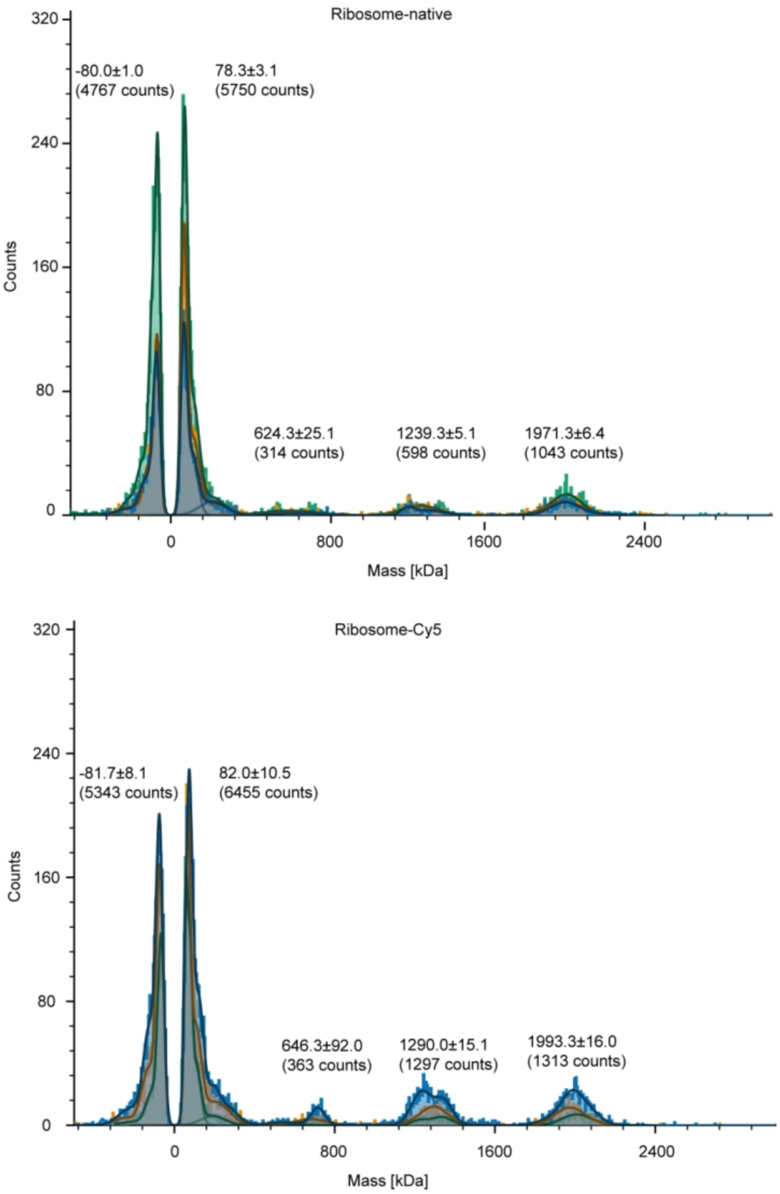


**Figure S12: Mass photometry data show ribosome-Cy5 was partially disassembled after fluorescence labelling.** The mass photometry experiments were calibrated with BSA/IgG/thyroglobulin and used to measure the relative mass of ribosome (top) and labeled ribosome (bottom). The peak of 70S (2 MDa) was visible in both cases, but the peak near 50S (1300kDa) and 30S (700kDa) was significantly increased in the sample containing the labeled ribosome. 3 repeats were performed and averaged.

### SI notes I: Ribosome partitioning in proteinosomes.

Mass photometry was used to confirm the size of ribosomes after fluorescent labeling. Mass photometry confirmed that the labeled ribosomes were truncated, with 30S, 50S fractions increased compared to unlabeled ones (**Figure S11**). Thus, the partitioning results provide information on the location of 30S and 50S subunits. Nevertheless, with the fluorescence microscopy, we observed restricted partitioning of ribosomes into cationized proteinosomes, which may explain the low expression level inside these proteinosomes. The native 23roteinosomes, however, have showed a mixed population of ribosome locating.

In addition, proteinosomes were incubated in a diluted PURExpress, then washed in buffer by centrifugation. These native proteinosomes, crosslinked with $BS(PEG)_{2000}$-crosslinked was imaged on TEM with negative staining (**Figure 1H**). Here, the ribosome was observed to be crowded inside the proteinosome lumen with some leakage, which indicates the partitioning of complete *E. Coli* 70S ribosome into the proteinosome, but also a possible physical crowding introduced by ribosome or other proteins. It was not possible to observe the ribosomes within the proteinosomes using the cross-sectioned proteinsosomes due to the chemical preparation steps which led to ribosome degradation and diffusion out of the proteinosome.

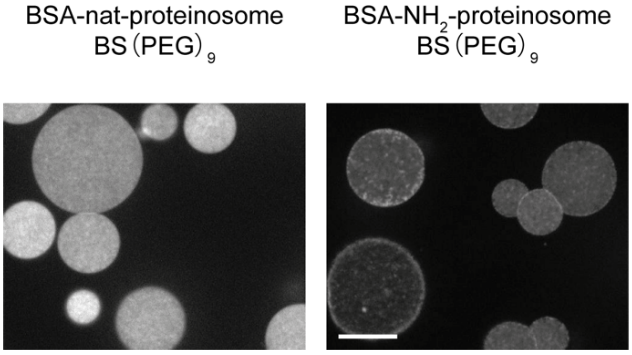


**Figure S13**: **Proteinosomes containing Plasmid DNA crosslinked with $BS(PEG)_9$.** Nat-BSA proteinosome and $BSA\text{-}NH_2$ proteinosome containing plasmid DNA were stained with 10 µM EvaGreen and imaged on a confocal spinning disk microscopy using a 60x/1.3 NA objective. Plasmid DNA signal were homogenously distributed inside native proteinosome (left) while accumulated on the outer layer in cationized proteinosomes (right).  Scale bar: 20 μm.

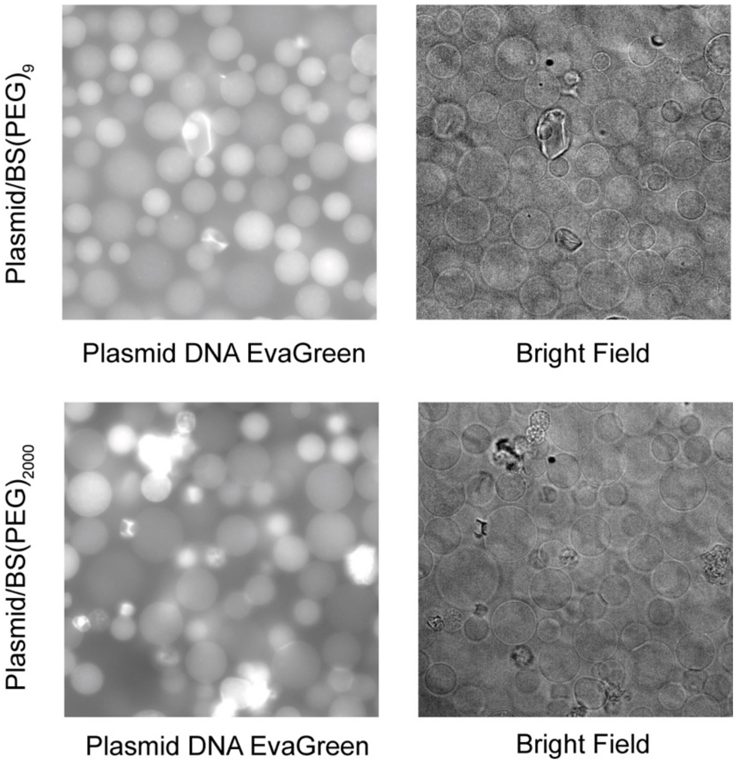


**Figure S14: Long term stability of the native proteinosomes.** Plasmid-DNA encapsulated proteinosomes were incubated in water for 11 months. Water was changed a few times during the storage period. The proteinosomes were mixed with EvaGreen and imaged with a fluorescence microscopy with 40x/0.8 NA objective. Results show intact proteinosomes after a storage period. Scale bar 20 μm.

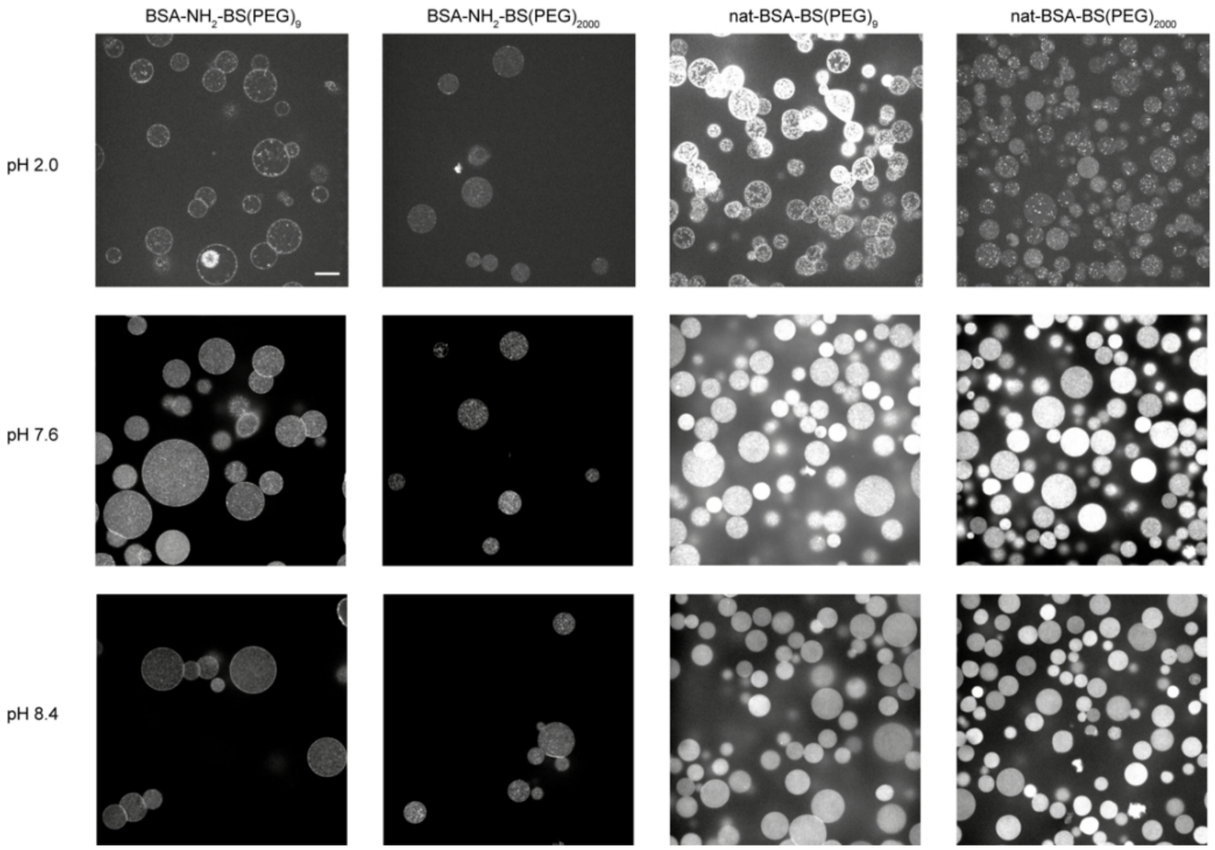


**Figure S15: Effect of pH on BSA proteinosomes containing DNA.** Cationized BSA proteinosome or native BSA proteinosomes were prepared by pre-loading plasmid and crosslinking with BS(PEG)$_9$ or BS(PEG)$_{2000}$. The proteinosomes were further incubated with EvaGreen to label the DNA in buffer at pH 2.0, 7.6 and 8.4. The samples were imaged at 37˚C on a spinning disk confocal microscope using a 60x/1.3 NA objective. Scale bar 20 µm.

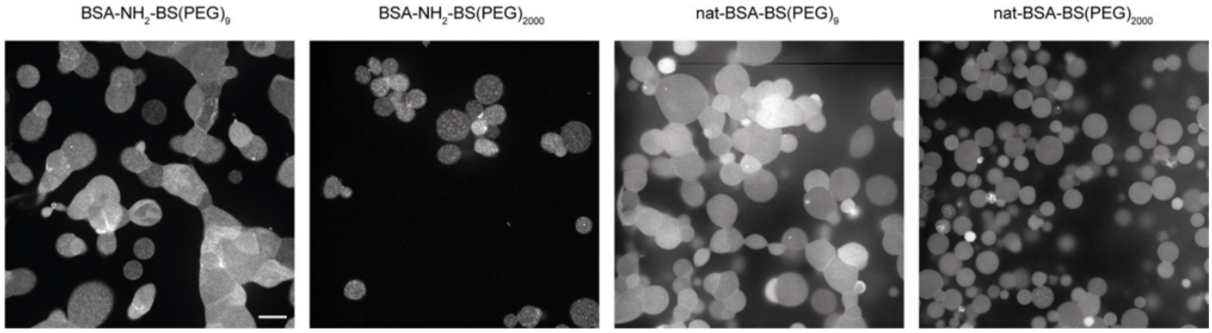


**Figure S16: Effects of osmolarity change on different BSA proteinosome containing plasmid DNA.** Cationized BSA proteinosome or native BSA proteinosomes with plasmid were incubated with 1µM EvaGreen in ribosome dilution buffer (pH 7.6) together with 1 M sucrose at 37˚C. The samples were imaged at 37˚C on a spinning disk confocal microscope using a 60x/1.3 NA objective. At high osmolarity, native proteinosomes with BS(PEG)$_{2000}$ were the most stable. Scale bar 20 μm.

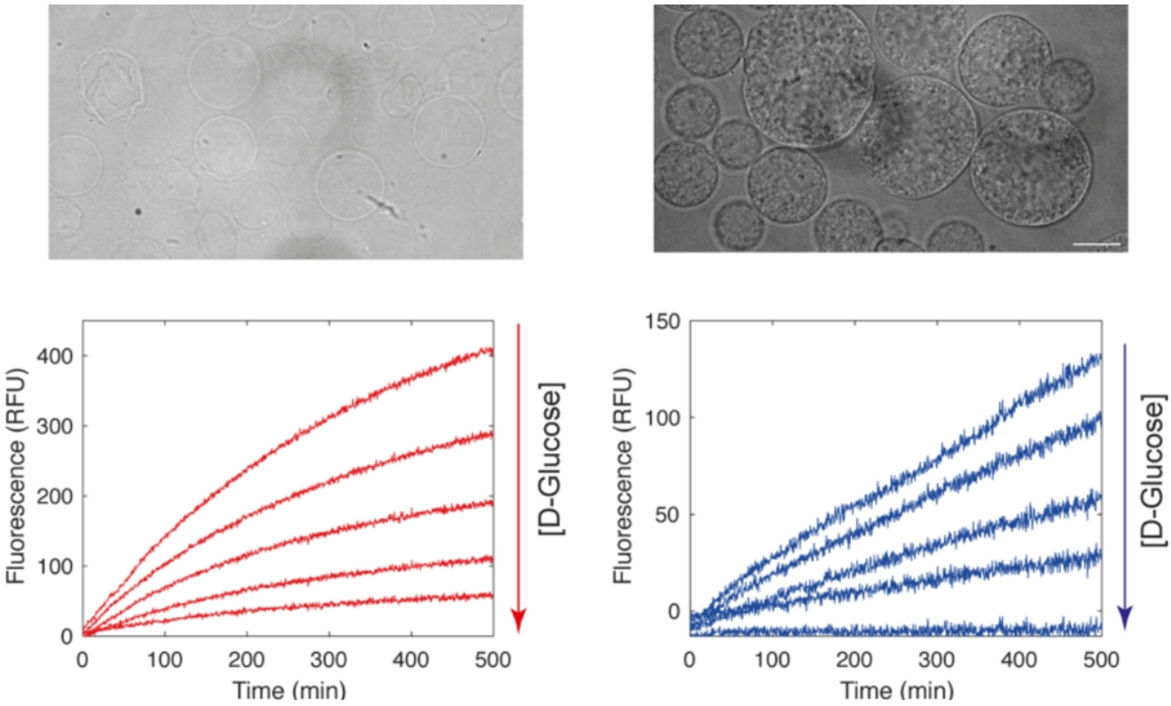


**Figure S17 Enzymatic activity of nat-GOx and GOx-NH$_2$ proteinosome.** Proteinosomes were imaged under a bright field microscopy (Top) after the GOx-glucose reaction. GOx-glucose-HRP activity was monitored *via* Amplex Red on a Tecan 20M plate reader with 0-50 μM D-Glucose (Bottom). The experiements were repeated 3 times with different batches of proteinosomes. However it remained difficult to quantify the protein density on the proteinosomes, thus here the kinetics was to illustrate the activity of both kind of proteinosomes. Scale bar 10 μm.

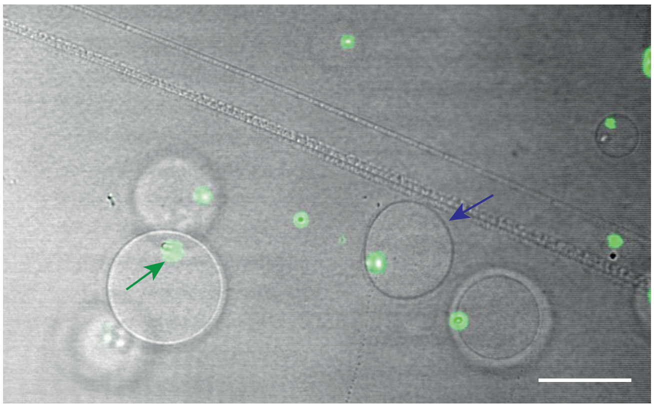

**Figure S18** PRM5/plasmid DNA phase separation (green arrow) inside nat-BSA proteinosome (blue arrow). DNA was stained with EvaGreen**.** As previously studied [9], PRM5 and negatively charged RNA or DNA phase-seperation at high concentration. Here, upon the adding of PRM5 into the DNA contained proteinosome, phase separation was observed inside the proteinosome. Scale bar: 20 μm.

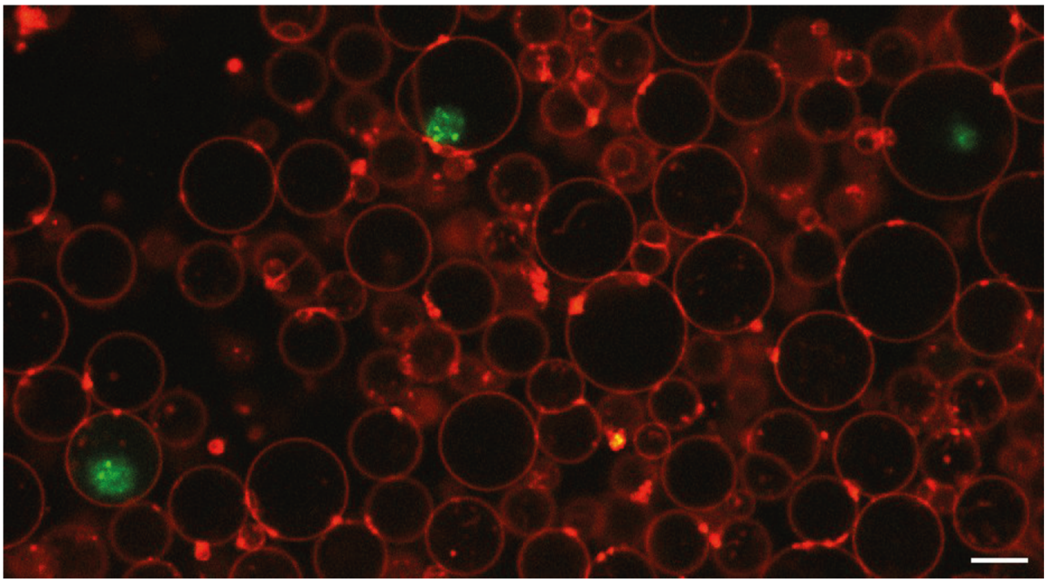

**Figure S19** Dextran-FITC (green, 2000 kDa) and mCherry-Plasmid DNA contained proteinosome in liposome. Plasmid contained proteinosomes were incubated in 1xPURExpress and the mixture was emulsified with egg PC/DiD (red) to generate liposome via centrifugation. The sample was incubated at 37˚C and imaged via a confocal microscope using a 63x/1.3 NA Plan-Apochromat objective. The result showed the encapsulation of proteinosome inside liposome in the buffer containing cell free machinaries. However, no protein expression was observed in 12 hours. Scalebar: 10 μm.

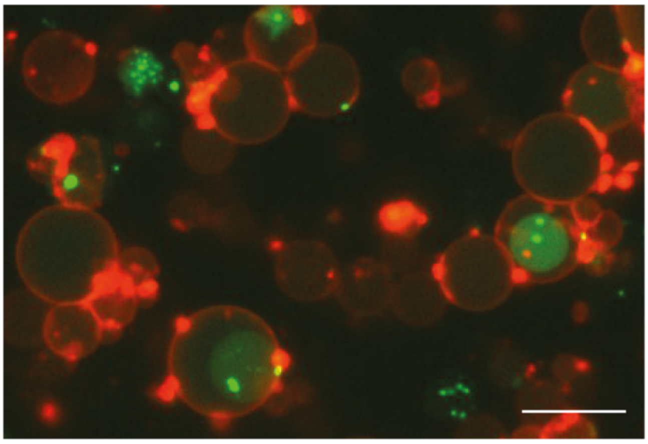

**Figure S20** PRM5-mCherry plasmid DNA contained proteinosome in liposome. Plasmid contained proteinosomes were incubated in PRM5 and Evagreen contained glucose solution. The mixture was emulsified with egg PC to generate liposome. DNA was stained in green while the liposomes were stained with DiD (red). As previously studied [9], PRM5 and negatively charged RNA or DNA phase-seperation at high concentration. Here we used the mechanism to achieve the phase separation in proteinosome inside the liposome. Scale bar: 15 μm.

**Movie S1: Cell free expression of mCherry in nat-BSA-BS(PEG)$_{2000}$ proteinosomes.** Transcription of mRNA (green, DHFBI stained) and translation of mCherry (red) were monitored at two different focal planes within the proteinosomes and on the surface of an agarose beads. The samples were imaged at 37˚C on a spinning disk confocal microscope using a 20x/0.5 NA objective.
